# A single *Citrobacter rodentium* infection in Pink1 knockout and wildtype mice leads to regional blood-brain-barrier perturbation and limited microglial activation without dopamine neuron axon terminal loss

**DOI:** 10.1101/2024.12.24.630165

**Authors:** Sriparna Mukherjee, Vladimir Grouza, Alex Tchung, Amandine Even, Moein Yaqubi, Marius Tuznik, Tyler Canon, Sherilyn Junelle Recinto, Christina Gavino, Marie-Josée Bourque, Nicolas Giguère, Heidi McBride, Michel Desjardins, Samantha Gruenheid, Jo Anne Stratton, David A. Rudko, Louis-Eric Trudeau

**Affiliations:** Department of pharmacology and physiology, Faculty of Medicine, Université de Montréal, Montreal, Quebec, Canada; Department of neurology and neurosurgery, Faculty of Medicine, McGill University, Montreal Neurological Institute, Montreal, Quebec, Canada; Department of microbiology and immunology, Faculty of Medicine, McGill University, Montreal, Quebec, Canada; Department of pathology and cell biology, Faculty of Medicine, Université de Montréal, Montreal, Quebec, Canada; Department of neurosciences, Faculty of Medicine, Université de Montréal, Montreal, Quebec, Canada; SNC and CIRCA research groups, Université de Montréal, Montreal, Quebec, Canada; Aligning Science Across Parkinson’s (ASAP) Collaborative Research Network, Chevy Chase, MD 20815

**Author notes:** **Address correspondence to: Dr. Louis-Eric Trudeau Professor**, Department of pharmacology and physiology Faculty of Medicine, Université de Montréal Montreal, Quebec, Canada. Denotes equal contribution (Vladimir Grouza, Alex Tchung, Amandine Even).

## Abstract

A growing body of research suggests a link between immune system activation and the development of Parkinson’s disease (PD). Previous work showed that repeated gastrointestinal infection with *Citrobacter rodentium* can induce PD-like motor dysfunction in *Pink1* knockout (KO) mice, along with immune cell infiltration into the brain. To better understand mechanisms underlying immune-mediated brain attack in this model, we tested whether mild infections are sufficient to increase blood-brain barrier (BBB) permeability and trigger brain inflammation. *Pink1* wild-type (WT) and KO mice were infected with *C. rodentium*, and gadolinium-enhanced magnetic resonance imaging (MRI) was performed at days 13 and 26 post-infection to assess BBB integrity. Quantitative MRI analysis revealed increased BBB permeability at day 26 in both WT and KO mice, particularly in the striatum, dentate gyrus, somatosensory cortex, and thalamus. Notably, this permeability was not associated with changes in tight junction protein expression or dopamine system markers in the striatum at either time point. However, persistent microglial activation was observed at day 26 post-infection, along with elevated levels of inflammatory mediators such as eotaxin, IFN-γ, CXCL9, IL-17, and MIP-2 in the striatum. Additionally, serum levels of IL-17 and CXCL1 were increased in infected *Pink1* KO mice. Flow cytometry revealed neutrophil infiltration in the brain at day 26 post-infection. Finally, a bulk RNA-seq transcriptome analysis revealed that gene sets related to synaptic function were particularly influenced by the infection and that inflammation-related genes were upregulated by the infection in the Pink1 KO mice. These findings support the hypothesis that even mild gastrointestinal infections can increase BBB permeability, disrupt brain homeostasis, and promote chronic neuroinflammation. In genetically susceptible individuals, such as those with *Pink1* deficiency, this may represent a first hit that contributes to subsequent induction of PD pathology with aging.

**Author summary:** We hypothesize that immune system activation is linked to the development of Parkinson’s disease (PD). Previous work revealed that repeated gastrointestinal infections with *Citrobacter rodentium* causes PD-like symptoms and immune cell invasion in the brain of *Pink1* knockout (KO) mice. In the current study, we tested whether a single mild gut infection alters blood-brain barrier (BBB) permeability and causes brain inflammation. We infected *Pink1* WT and KO mice with *Citrobacter rodentium* and used gadolinium-enhanced MRI to detect BBB permeability changes at 13- and 26-days post-infection. Results showed increased BBB permeability in specific brain regions at 26 days. While tight-junction and dopamine (DA)-related proteins remained unchanged, we observed altered expression of synaptic genes, chronic microglial activation, elevated inflammatory markers, and neutrophil infiltration in the brain. Our findings suggest that even mild gastrointestinal infections can increase BBB permeability, potentially enabling immune cell infiltration into the brain and exacerbating pathways implicated in the development of PD, particularly among individuals with genetic risk factors.

## Introduction

Parkinson’s disease (PD) is one of the fastest-growing neurological disorders, largely driven by population aging, as advancing age is a significant risk factor for the condition (1, 2). Although considered for decades as a selective disease of neurons, there is growing recognition that PD is a multi- system disorder involving not only the loss of DA neurons in the ventral midbrain, but also the degeneration of peripheral innervation, mitochondrial dysfunction in multiple cell types and a disruption of the gut microbiome. Based on the presence of Lewy pathology in multiple regions of the body at different time points after the diagnosis of PD, it has been speculated that PD pathology begins in the enteric nervous system or in the olfactory system before spreading within the central nervous system, commonly known as the ‘body-first’ hypothesis (3, 4). The protein α-synuclein (α-syn), a major constituent of Lewy bodies in the brain, has also been reported to accumulate in the gastrointestinal tract, heart and skin, providing further indirect evidence for the implication of system-wide perturbations in PD (5).

Several studies have reported significant differences in the composition of the gut microbiota between PD patients and healthy controls. For example, a reduction in the abundance of *Prevotellaceae* and an increase in *Enterobacteriaceae* have been observed in PD patients, which were positively correlated to the severity of postural instability (6). Gut dysbiosis in PD could perhaps lead to increased peripheral inflammation secondary to changes in gut microbiome-derived short chain fatty acids, which can promote elevated intestinal permeability commonly referred to as “leaky gut”. This physiologic condition can trigger systemic inflammation and possibly contributes to neuroinflammation observed in PD (7). Bacterial infections are otherwise progressively being recognized as a possible trigger for the onset of PD. *Helicobacter pylori* infection is prevalent among patients with PD (8). Studies have consistently shown that individuals with *H. pylori*-related ulcers have a higher risk of developing PD compared to healthy individuals of a similar age (9). Additionally, *Mycobacterium tuberculosis*, *Porphyromonas gingivitis* and *Clostridium difficile* infections have been identified to be risk factors for the onset of PD (10).

Although most cases of PD are considered sporadic, familial forms of the disease occur due to mutations in a growing list of genes including *Pink1*, *Parkin*, *LRRK2*, *SNCA*, *DJ-1*, *GBA1* and *ATP13A2*. Interestingly, most mouse models carrying mutations or deletions of these genes do not develop PD-like symptoms. Such observations are compatible with the hypothesis that genetic vulnerability factors need to interact with other precipitating factors to lead to brain or peripheral pathology. Environmental factors including infections and changes in gut microbiome can be hypothesized to represent such a second hit, leading to PD. The collaborative interplay of these factors for the development of PD has been previously studied in animal models in which pesticides, viral or bacterial infections have been shown to act as disease initiators. As an example, the *E coli* protein curli was shown to facilitate α-syn aggregation in brain followed by microglial activation and motor deficits in a mouse model (11). In another study, *Proteus mirabilis* administration with MPTP in mice led to PD-like motor symptoms and DA neuron damage (12). The bacterial infection in this model was shown to weaken the BBB and to increase the quantities of α-syn fibrils in both the brain and colon of these mice. Bacterial infections could lead to the development of PD-like pathology through immune system activation, as revealed by recent work showing that four cycles of gastrointestinal infection with *Citrobacter rodentium* triggers autoimmune-like mechanisms in Pink1 KO mice and the generation of mitochondria specific CD8+ T cells in the periphery and CNS (13). Motor impairment and loss of striatal DA neuron terminals were also observed in this animal model at 6 months after infections. These findings reinforce the hypothesis that immune mechanisms play a key role in the development of PD and that loss of function of proteins such as Pink1 can enhance innate and adaptive immune responses triggered by pathogens. Once triggered, the increased abundance of autoreactive CD8+ T cells may be sufficient to induce DA system dysfunction in both Pink1 KO and WT mice, as revealed in a recent study using an adoptive transfer strategy (14). Although the previous *Citrobacter rodentium* study clearly illustrates the long-term outcome of intestinal infection, it is unclear whether single infections are sufficient to perturb brain homeostasis and immune cell infiltration in the brain.

In the present work, we therefore explored whether a single gastrointestinal *Citrobacter rodentium* infection in Pink1 WT and KO mice can perturb the BBB and initiate inflammatory signals in the brain. We found that at 26 days after the infection, although the integrity of the DA system in the brain is not yet compromised, changes in BBB permeability occur both in Pink1 WT and KO mice and microglial activation as well as elevation of pro-inflammatory cyto-chemokines are detected.

## Results

### *Citrobacter rodentium* infection causes splenomegaly and increases colon weight in Pink1 WT and KO mice

Our initial objective was to test the hypothesis that bacterial infection of the gastrointestinal tract can cause changes in brain permeability. To achieve this, we first validated the effectiveness of *Citrobacter rodentium* infection in DATcre;Ai9;Pink1 transgenic mice (thereafter called DATcrePink1 mice). Considering that gastrointestinal *Citrobacter* infection leads to a peak of infection at about 10-14 days after gavage and resolves thereafter (13), we harvested the tissues at either day 13 (infection peak) or day 26 (infection resolved) post-infection (Fig 1A). A stock of chloramphenicol-resistant DBS100 strain was grown overnight and the next day, 200μl of bacterial culture was used for oral gavage in both Pink1 WT and KO mice, while the control mice received an equal volume of the same culture broth used to grow the bacteria. At day 6 post infection, feces were collected from the infected mice. Analysis of three independent cohorts showed that the feces contained 10^7^-10^10^ CFU of *Citrobacter*/g in both Pink1 WT and KO mice (Fig1B), confirming an efficient intestinal infection in both genotypes. Also validating an effective infection, we observed a significant increase in the spleen index in both the Pink1 WT and KO infected mice as compared to the controls. The enlargement in spleen was higher at day 13 post infection as compared to day 26 (Fig 1C), in keeping with the fact that the infection is mostly resolved at day 26. Similarly, we found a higher colon index in the infected mice of both genotypes at day 13 post infection as compared to the controls, which persisted in the Pink1 KO mice at day 26 but not in the WT infected mice (Fig 1D). These findings suggest that there might be additional immune mechanisms at play in the Pink1 KO mice in response to the infection, leading to longer-lasting alterations.

**Fig 1.**
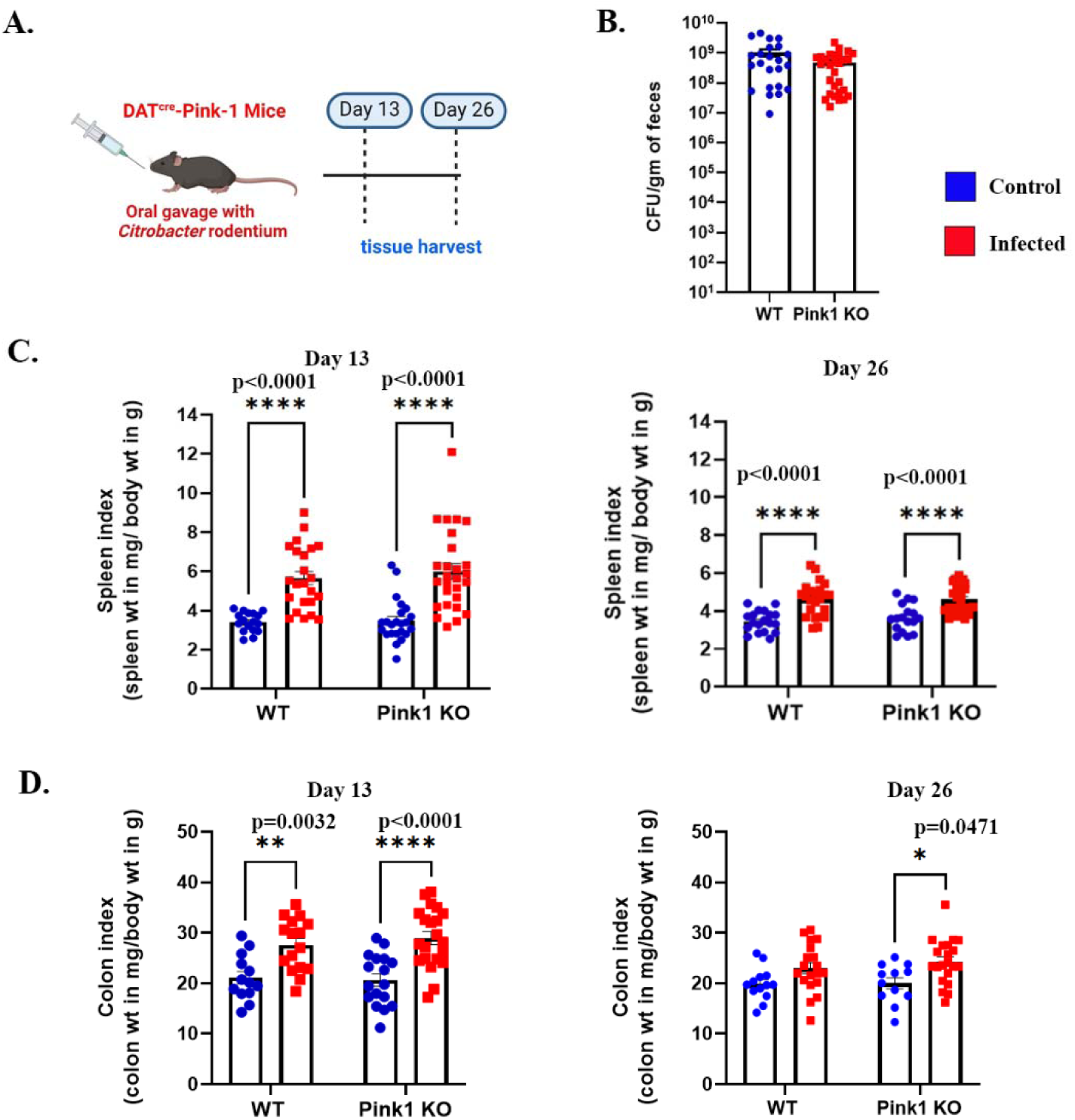
Infection with *Citrobacter rodentium* leads to splenomegaly and colon enlargement in both wild-type (WT) and PINK1 knockout (KO) mice. WT and Pink1 KO mice were infected with *C. rodentium* and at day 13 and day 26 post infection, tissue samples were collected. Created in BioRender. Mukherjee, S. (2026)https://app.biorender.com/citation/69824cdcd69996e0da8584e7. (A). 10^7-10^10 CFU of Citrobacter /g of feces in WT and Pink1 mice signifies successful intestinal infection in both genotypes. (B). Higher spleen index was noted in both WT and KO infected mice at day13 and day 26 post infection (C). Higher colon index in the infected mice of each genotype was observed only at day 13, while at day 26, colon enlargement persisted only in the infected KO mice (D). Data is representative of three independent experiments and presented as mean ± S.E.M. p value was determined by two-way ANOVA at 95% confidence and Tukey’s post hoc test.

### *Citrobacter rodentium* infection leads to a regional increase in blood brain barrier permeability in both Pink1 WT and KO mice

MRI-based T_1_ mapping, combined with the intravenous administration of gadolinium (0.4mmol/kg) as a contrast agent, was used to examine the permeability of the BBB in Pink1 WT and KO mice. Imaging was performed using a 7 Tesla pre-clinical MRI system with a high amplitude gradient insert. The image acquisition and analysis pipeline are illustrated in Fig 2A-C. Successful delivery of the contrast agent was evidenced qualitatively by tissue hyperintensity in post-contrast magnitude images after delivery of gadolinium (Fig 2A). Image processing was performed by brain masking with the SHERM tool and registration of the template-based regions of interest (ROI) atlas (Fig 2B). A monoexponential signal model was used for T_1_ calculation, based on the data collected from VTR imaging pulse sequence (Fig 2B). An example of T_1_ maps measured pre and post CA delivery is shown in Fig 2C. Contrast agent delivery to the brain parenchyma was evidenced by a T_1_ shift to shorter values (Fig 2Cii). The degree of shift was quantified by calculating the Earth Mover’s Distance (EMD). T_1_ values were evaluated in specific anatomical locations of the brains of WT and Pink1 KO mice. Both infected and non-infected image data sets are represented in Supplementary Fig. 1A-D. Seven to 10 mice per genotype and infection were imaged days 13 or 26 after the infection. Quantification of T_1_ shortening in each brain ROI at day 13 post infection did not reveal any significant effect of the infection on the T_1_ relaxation constant (Fig 2D). However, there was a significant global difference between the two genotypes in the striatum, thalamus, dentate gyrus and primary somatosensory cortex, with WT mice showing larger T_1_ shortenings compared to the KO mice, suggesting possible baseline differences in BBB permeability between Pink1 WT and KO mice (Fig 2D). Importantly, at 26 days post infection, T_1_ shortenings were significantly larger in the infected mice for both Pink1 WT and KO mice in these same brain regions compared to th controls (Fig 2E). Our results demonstrate that a single intestinal infection can indeed trigger BBB disruption in the infected mice irrespective of genotype.

**Fig 2.**
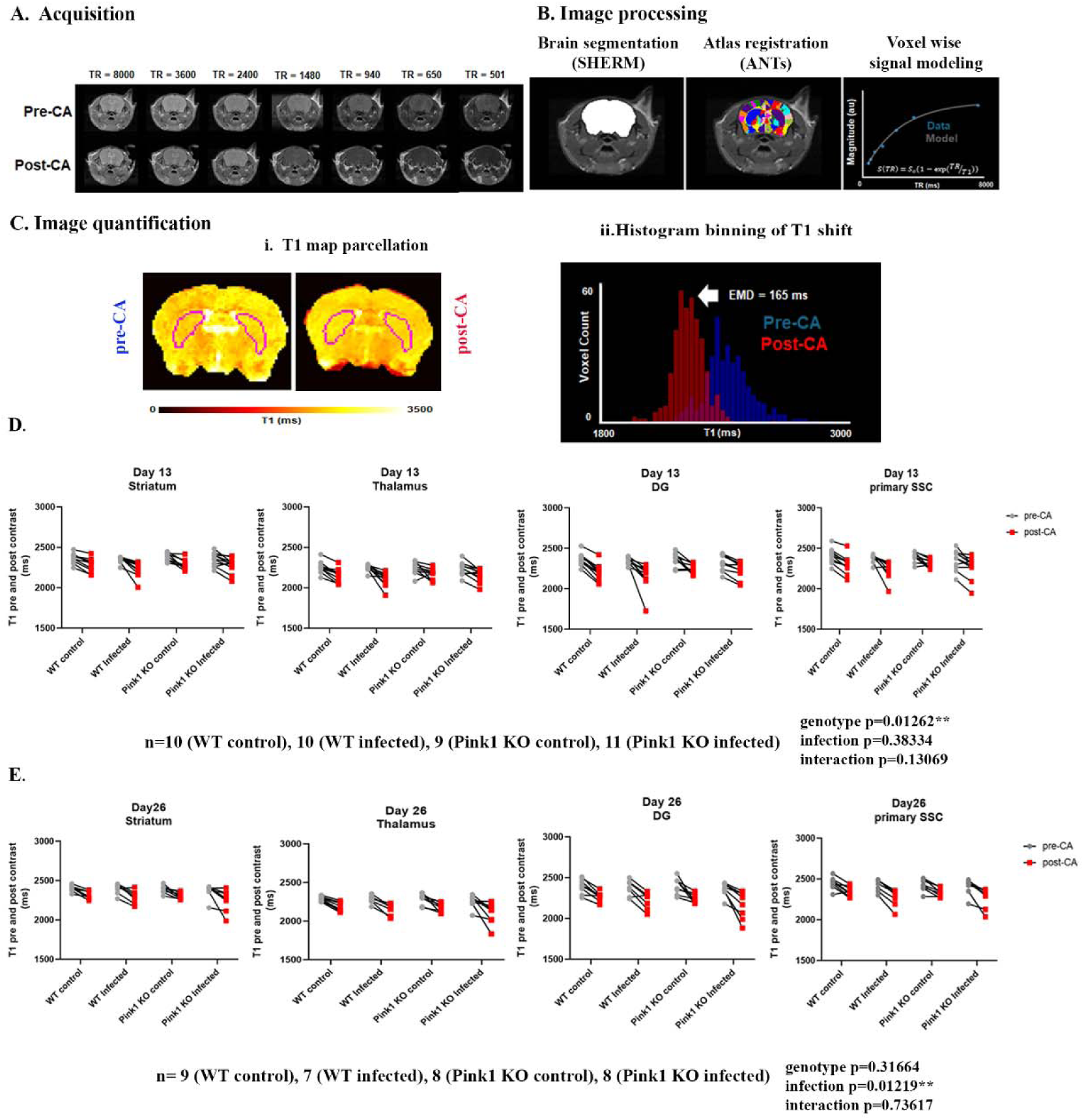
Illustration of image acquisition, processing pipeline and T1 analysis for assessment of regional BBB permeability with gadolinium-enhanced MRI in *Citrobacter* infected mice. MR images acquired with the RARE-VTR protocol are shown in (A). Successful delivery of th gadolinium-based contrast agent is indicated by the increased soft tissue hyperintensity in the shorter (<1480 ms) TR post-contrast agent (CA) images. The image processing and signal modeling of the RARE-VTR data are depicted in (B), including a pre-CA image. Brain masking accuracy with SHERM, registration of the template-based ROI atlas, and signal modeling were performed for each mouse (both non-infected and infected, across genotypes). Examples of T1 maps pre- and post-CA delivery, with the striatum ROI highlighted in magenta, are shown in (Ci). Gadolinium-based contrast agent (GBCA) infiltration into brain parenchyma was evidenced by the reduction in T1 values. The extent of this shift was quantified for each experimental mouse using Earth Mover’s Distance (C ii). The T1 signal shift in the striatum, thalamus, dentate gyrus, and primary somatosensory cortex at day 13 post-infection for both non-infected and infected mice of wild-type (WT) and Pink1 knockout (KO) genotypes is shown in (D). Similarly, the T1 signal changes pre- and post-CA for non-infected and infected mice of both genotypes at day 26 post-infection are presented in (E) [red: post contrast, grey: pre contrast].

### Expression of endothelial tight junction proteins is unchanged after infection

An increase in BBB permeability after infection could result from multiple mechanisms. One of these could be a reduction in the expression of endothelial tight junction proteins. To examine this possibility, we performed western-blot analysis from striatal lysates to examine the levels of tight junction proteins. The results did not reveal any significant changes in the levels of the endothelial tight junction proteins ZO-1 or ZO-2 at 13 or 26 days after the infection. On day 13, a significant decrease in ZO-1 was detected in the Pinkl KO mice compared to WT (Fig 3A-3B). At day 26, the expression of ZO-1 was modestly higher in the Pink1 KO mice compared to the WT, while the expression of ZO-2 was modestly elevated in the KO infected mice as compared to the WT infected mice (Fig 3C-3D). Globally these findings do not support the hypothesis that the increase in BBB permeability observed after infection was due to reduced expression of tight junction proteins.

**Fig 3.**
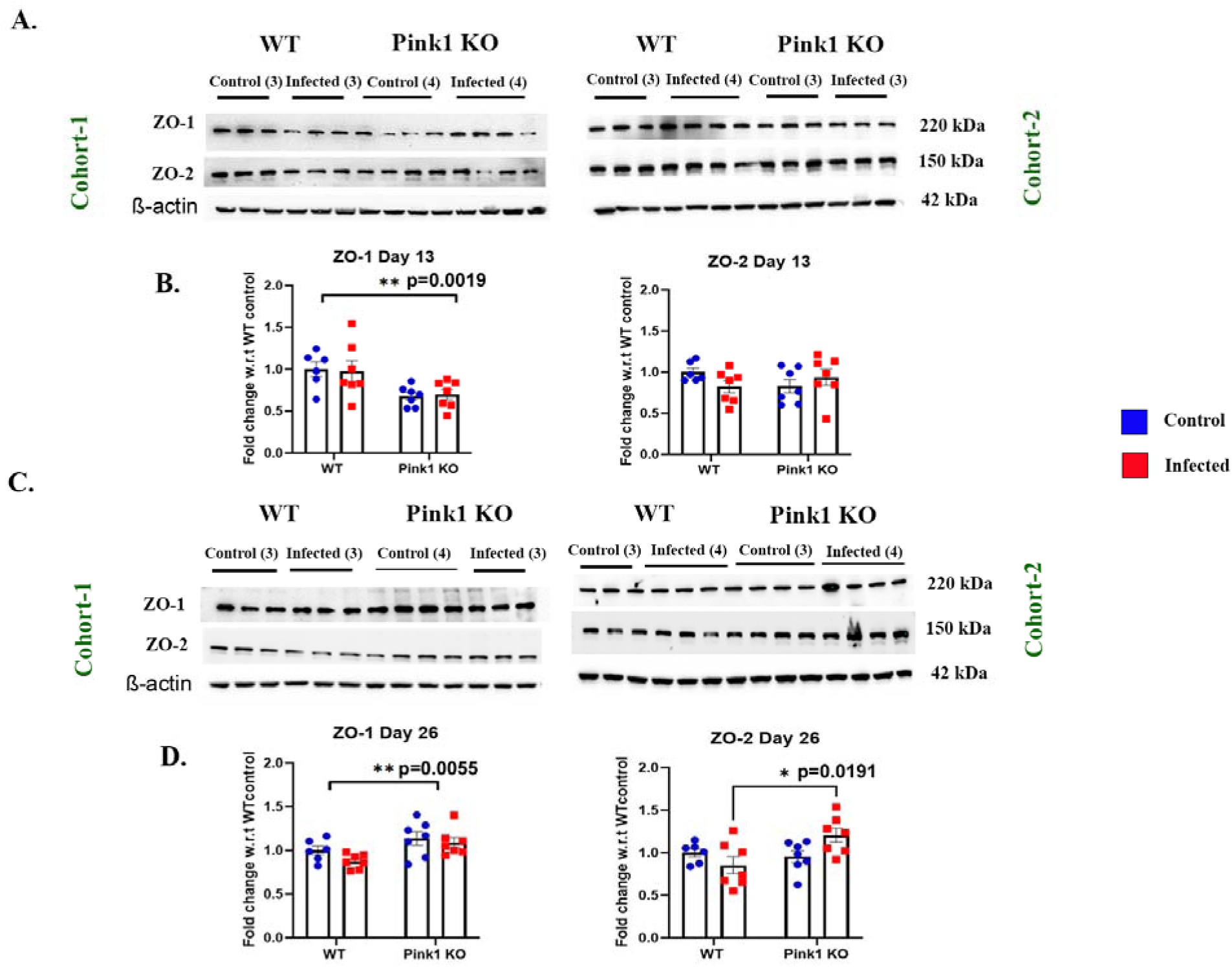
*Citrobacter rodentium* infection does not result in a reduction of endothelial tight junction proteins. Striatal lysates from the brain tissue of non-infected and infected mice were prepared, quantified and separated on SDS-PAGE and probed for ZO-1 and ZO-2. Expression of ZO-1 and ZO-2 at day 13 post infection is depicted in panels A and B, whereas the expression of these proteins at day 26 post infection is shown in panels C and D. Data is representative of two independent mouse cohorts and presented as mean ± S.E.M. p values were determined by twoway ANOVA at 95% confidence and Tukey’s post hoc test.

### *Citrobacter* infection is associated with an elevation of cytokines and chemokines in the striatum and periphery

As endothelial cell functions including tight junction functions can be regulated by a range of cytokines and chemokines (15, 16), it is possible that the changes in BBB permeability we detected after infection resulted directly or indirectly from local changes in the expression of such signaling molecules. To examine this, we evaluated the striatal cyto-chemokine profile at days 13 and 26 after the infection. At day 13, we noted significantly elevated eotaxin, IFN-γ, CXCL9 and MIP-2 in the infected mice compared to non-infected controls, irrespective of genotype (Fig 4A, 4B, 4C, 4E). We also detected an increase in the levels of IL-17 selectively in the Pink1 KO mice (Fig 4D). VEGF levels showed a decrease at day 13 in both genotypes (Fig 4F). These changes were temporally associated with the peak of infection as no significant differences remained at day 26 post infection. Such signals in the striatum could contribute to the creation of a microenvironment facilitating microglial activation or attracting peripheral immune cells to infiltrate in the CNS.

**Fig 4.**
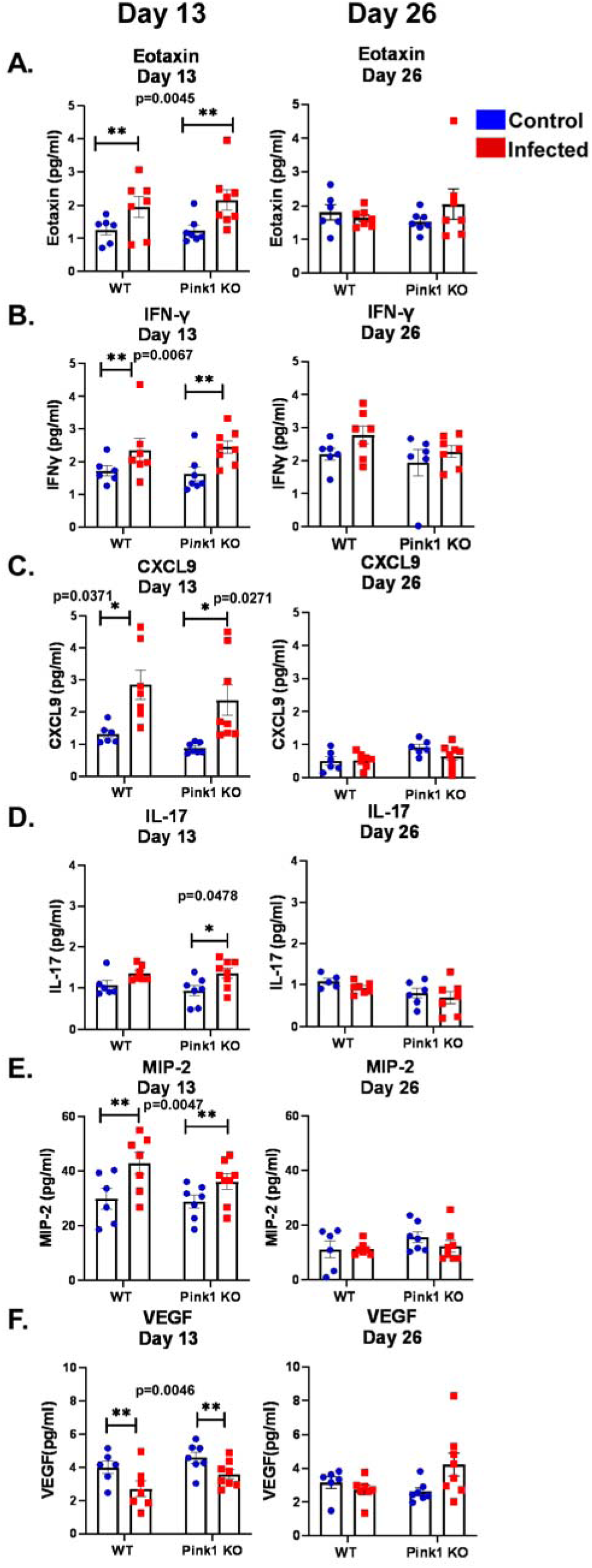
An increase in pro-inflammatory cyto-chemokines is noted in the striatum of *Citrobacter rodentium* infected wild-type (WT) and PINK1 knockout (KO) mice. Pro-inflammatory cytokines and chemokines were analysed in striatal lysates prepared from noninfected and infected WT and KO mice at day 13 and day 26 post infection. Alteration of Eotaxin (A), IFN-γ (B), CXCL9 (C), IL-17 (D), MIP-2 (E) and VEGF (F) are shown for both day 13 (left panels) and day 26 (right panels). Data is representative of three independent experiments and presented as mean ± S.E.M. p value were determined by two-way ANOVA at 95% confidence and Tukey’s post hoc test.

It may be speculated that the elevation of cytokines and chemokines detected here in the striatum at the peak of infection results from signals derived from peripheral circulating cytokines. To examine this, we collected serum samples from a subset of control and infected mice of both genotypes at 6, 13 and 26 days after the infection and quantified the levels of cytokines and chemokines (Fig 5A-5F). Although changes were limited at day 6, with only KC (CXCL1) being significantly elevated after the infection in both genotypes (Fig 5E, left panel), we detected a significant elevation of both IL-17 (p=0.0487) and IP-10 (p=0.0128) at day 13, here again in both Pink1 WT and KO mice (Fig 5B-5C, middle panel). Finally, on day 26, we detected a significant elevation of IL-17 (p=0.0021) and KC (CXCL1) (p=0.0158), this time only in Pink1 KO mice (Fig 5B and 5E, right panel). The results confirm the expected development of systemic inflammation after the infection. The temporal correlation of these changes with the elevation of striatal cytokines and chemokines also argues for a link between the two events. The observation of sustained changes in cytokines and chemokines in Pink1 KO mice further argues for the existence of enduring changes in inflammatory signals, especially in the KO mice.

**Fig 5.**
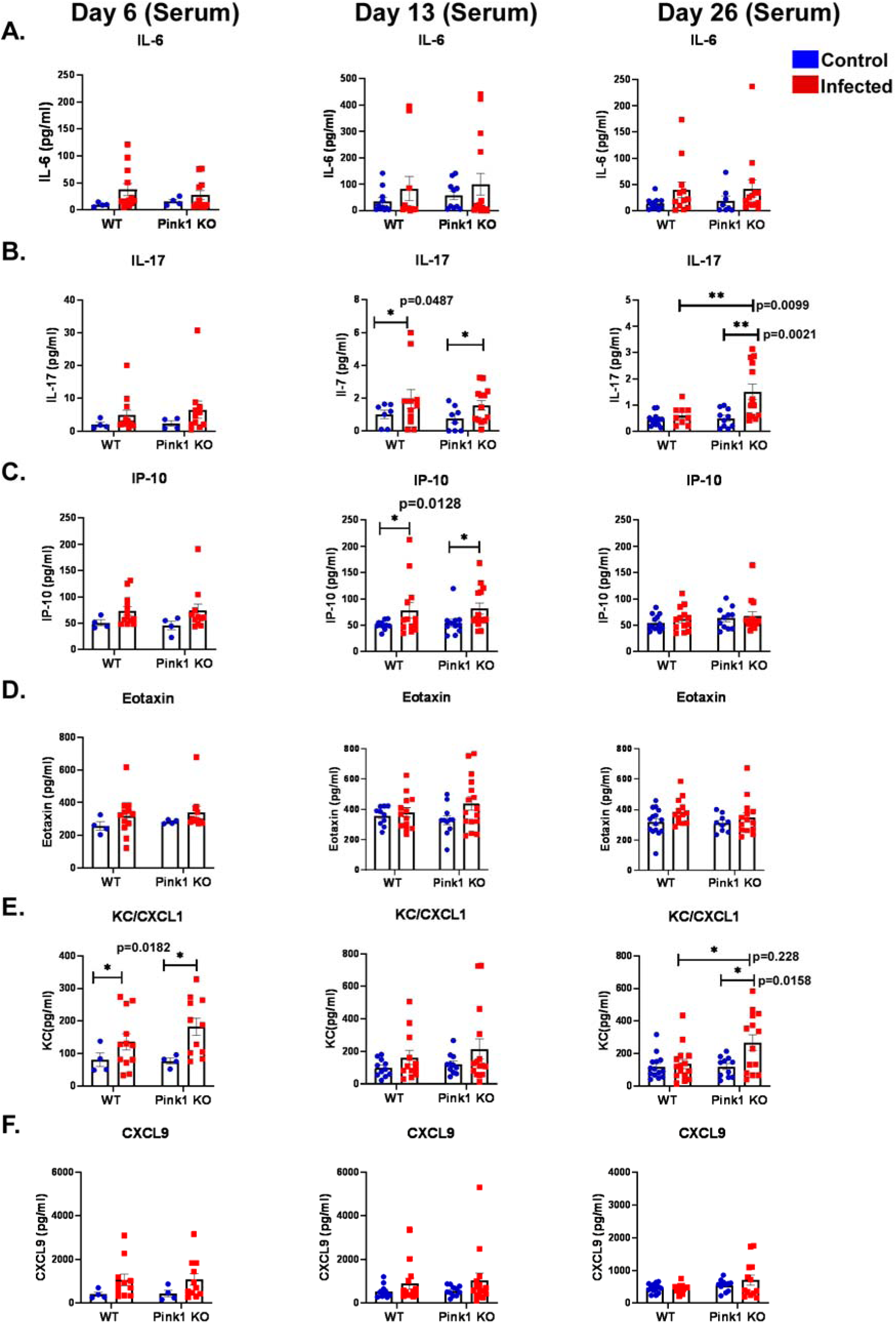
*Citrobacter rodentium* infection causes an increase in systemic pro-inflammatory cytokines and chemokines in wild-type (WT) and PINK1 knockout (KO) mice. Serum samples were collected from non-infected and infected mice of both genotypes at day 6, day 13 and day 26 post infection for the analysis of pro-inflammatory cytokines and chemokines. Data for IL-6 (A), IL-17 (B), IP-10 (C), Eotaxin (D), KC/CXCL1 (E) and CXCL9 (F) are shown for day 6 (left panels), day 13 (middle panels) and day 26 (right panels). Data is representative of three independent experiments and presented as mean ± S.E.M. p value were determined by two-way ANOVA at 95% confidence and Tukey’s post hoc test.

### Dopamine neuron terminals are unaffected by a single *Citrobacter* infection

The increase in BBB permeability and accompanying changes in cytokines and chemokines could potentially have a negative impact of the integrity of the brain DA system. The axonal domain of these neurons has been hypothesized to be particularly vulnerable to inflammatory and other neurotoxic signals (17–19). To examine this, the expression of tyrosine hydroxylase (TH) and of the DA transporter (DAT) was quantified by western blot from striatal lysates. We detected no significant changes in total striatal TH and DAT levels at day 13 (Fig 6A-6B). These results were validated by quantitative immunohistochemistry (Fig 6C) and similarly, we detected no significant changes in the intensity or surface area of TH or DAT in the dorsal or ventral striatum (Fig 6D-6E). Likewise, at day 26, no detectable changes in the levels of TH or DAT were detected in response to the infection (Fig 7A-7E), except for a surprising increase in total DAT levels in the western blot experiments (Fig. 7B), something that was not validated in the immunohistochemistry experiments (Fig 7D-7E). Our results suggest that the inflammatory and other potentially neurotoxic signals generated following a single gut infection are not sufficient to compromise the structural integrity of the DA system at the time points examined.

**Fig 6.**
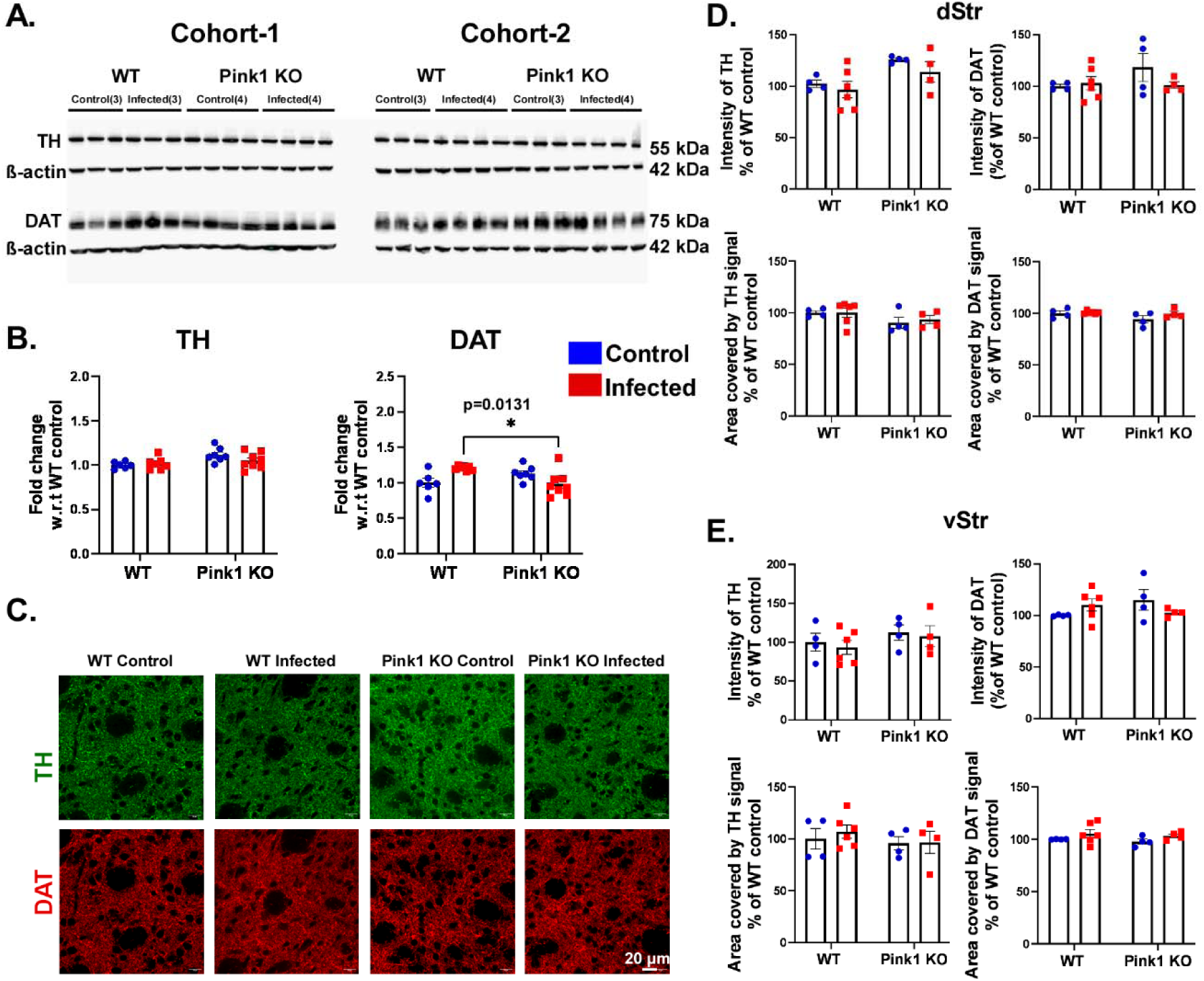
*Citrobacter rodentium* infection does not impact the structural integrity of dopamine neuron terminals in wild-type (WT) and PINK1 knockout (KO) mice at day 13 post infection. Striatal lysates were subjected to western blot and probed for tyrosine hydroxylase (TH) and dopamine transporter (DAT) (A). The blots were quantified to measure the change in expression of TH and DAT among the non-infected and infected experimental groups of both genotypes (B). The same protein expression in striatum was also analysed by immunofluorescence staining and quantification. Representative confocal striatal images are shown in (C), scale bar 20μm. Quantification of the TH and DAT intensity as well as the area covered by their signal were performed for both dorsal (D) and ventral striatum respectively (E). Data is representative of two independent mouse cohorts and presented as mean ± S.E.M. p value were determined by twoway ANOVA at 95% confidence and Tukey’s post hoc test.

**Fig 7.**
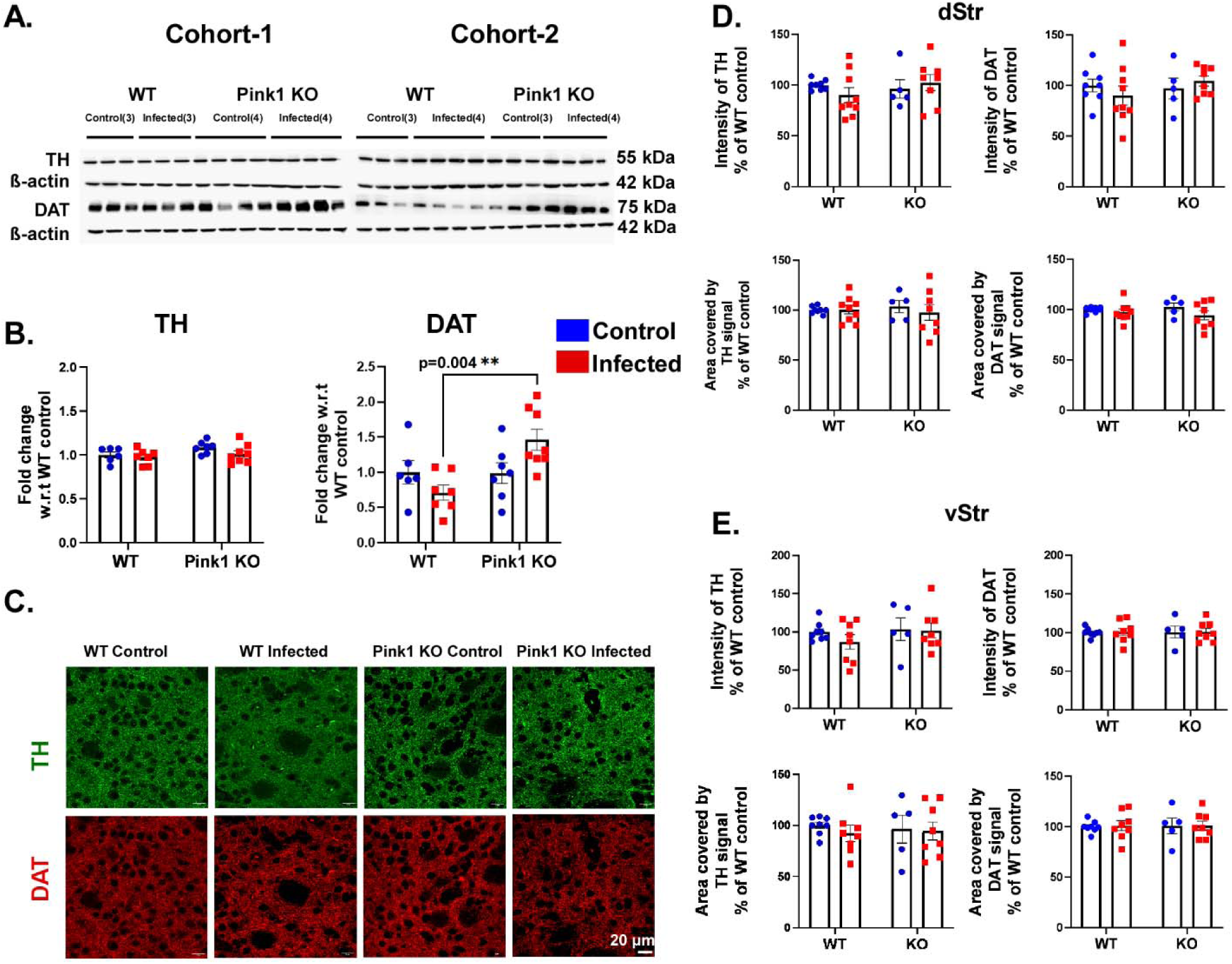
*Citrobacter rodentium* infection does not affect the structural integrity of dopamine neuron terminals in wild-type (WT) and PINK1 knockout (KO) mice at day 26 post-infection. Striatal lysates were analyzed by western blot, probing for tyrosine hydroxylase (TH) and dopamine transporter (DAT) (A). The blots were quantified to assess changes in TH and DAT expression between non-infected and infected groups across both genotypes (B). Protein expression in the striatum was also evaluated through immunofluorescence staining and quantification. Representative confocal images of the striatum are shown in (C). The intensity and area of TH and DAT signals were quantified in both the dorsal (D) and ventral striatum (E). Data represent two independent cohorts of mice and are presented as mean ± S.E.M. Statistical significance was determined using two-way ANOVA with Tukey’s post hoc test at a 95% confidence level.

### Quantitative analysis reveals preserved dopamine neuron populations in the SNc and VTA

To determine whether *Citrobacter rodentium* infection induced dopaminergic neurodegeneration, we estimated the quantity of tdTomato positive neurons in the ventral tegmental area (VTA) and substantia nigra pars compacta (SNc) using automated image analysis. Large-area z-stack images were acquired and processed with extended depth of field reconstruction, followed by machine learning-based detection of tdTomato positive neurons within anatomically defined regions of interest corresponding to the SNc and VTA.

Across both regions, the number of tdTomato positive neurons was comparable between control and infected mice in both WT and Pink1 knockout groups (Fig 8A-8C). No statistically significant differences were observed between treatment conditions or genotypes. These data indicate that infection does not result in detectable DA neuron loss in the SNc or VTA at day 26 post infection.

**Fig 8.**
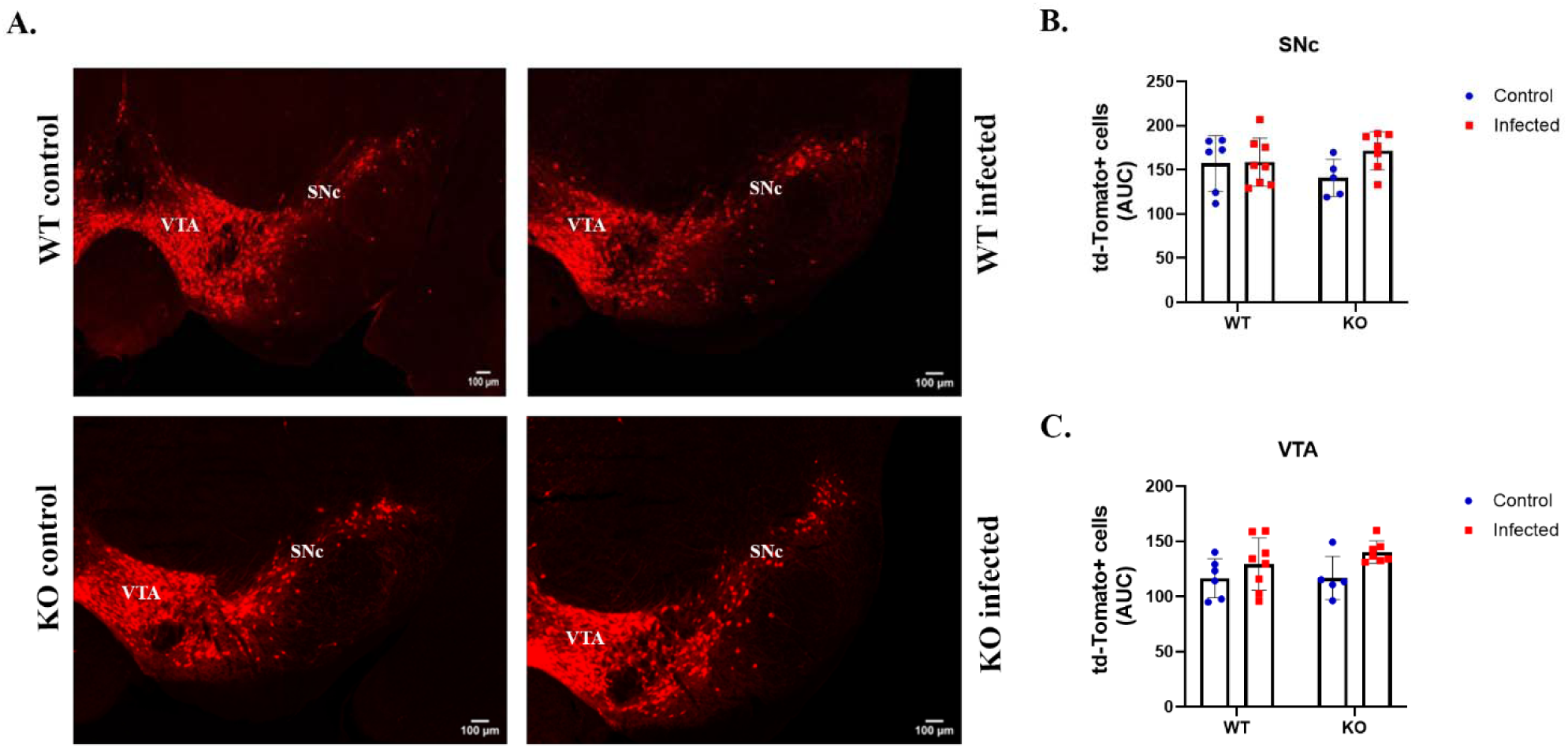
*Citrobacter rodentium* infection does not alter the number of dopamine neurons in the mesencephalon. 40 μm thick mesencephalic brain slices were processed for RFP immunofluorescence to visualize tdTomato-expressing DA neurons (A). The tdTomato-positive neurons were detected using a machine learning-based algorithm on each slice within the substantia nigra pars compacta (SNc) (B) and ventral tegmental area (VTA) (C). For each animal, the number of cells for each slice was plotted based on its corresponding bregma rostro-caudal coordinates and the estimated total number of cells was represented using the area under the obtained curve (AUC). Data are presented as mean ± SEM. Statistical significance was assessed using two-way ANOVA followed by Tukey’s post hoc test at a 95% confidence level. No statistically significant differences (p > 0.05) were detected among groups.

### Microglial but not astrocytic alterations are observed in the striatum after infection

Although the integrity of the DA system was not compromised after a single *Citrobacter* infection, it is possible that such infections initiate neuroinflammatory cascades in the brain that could set the stage for subsequent environmental signals, such as additional bacterial or viral infections, to promote the development of PD pathophysiology. Considering the broad literature showing that systemic inflammatory signals can promote glial activation in the brain and promote the establishment of neurotoxic cascades, we examined the state of microglia and astrocytes in the striatum. At day 13 post infection, the microglial marker IBA1 was found to be significantly up-regulated in the WT infected mice and downregulated in the KO infected mice, suggesting that the infection led to the activation of different signalling cascades in Pink1 WT and KO mice (Fig 9A-9B). At this same time point, the expression of the astrocyte marker GFAP was unaltered (Fig 9 A-9B). Because the shape of microglia has been demonstrated to correlate with different stages of their activation state, we also examined striatal sections by immunohistochemistry, here again quantifying IBA1 immunoreactivity (Fig 9C). A detailed morphological analysis failed to detect any significant changes in the overall density of IBA1-positive cells (Fig 9D), the size of their soma (Fig 9E), the intensity of IBA1 in the soma (Fig 9F), the circularity of the soma (indicative of the polarization of the membrane) (Fig 9G) or the area covered by microglial processes (Fig 9H). Interestingly, at day 26 post infection, a significant elevation in IBA1 levels in the striatum was selectively observed in the Pink1 KO infected mice, suggestive of a long-lasting state of chronic inflammation in the striatum of the Pink1 KO mice (Fig 10A-10B). Similarly to day 13, we did not observe any significant changes in GFAP expression (Fig 10A-10B) and immunohistochemical observations of IBA1 failed to reveal any changes in microglial morphology (Fig 10C-10H). These results argue for the establishment of brain inflammation implicating microglial cells and suggest the possibility of longer-lasting changes in the Pink1 KO mice compared to the WT.

**Fig 9.**
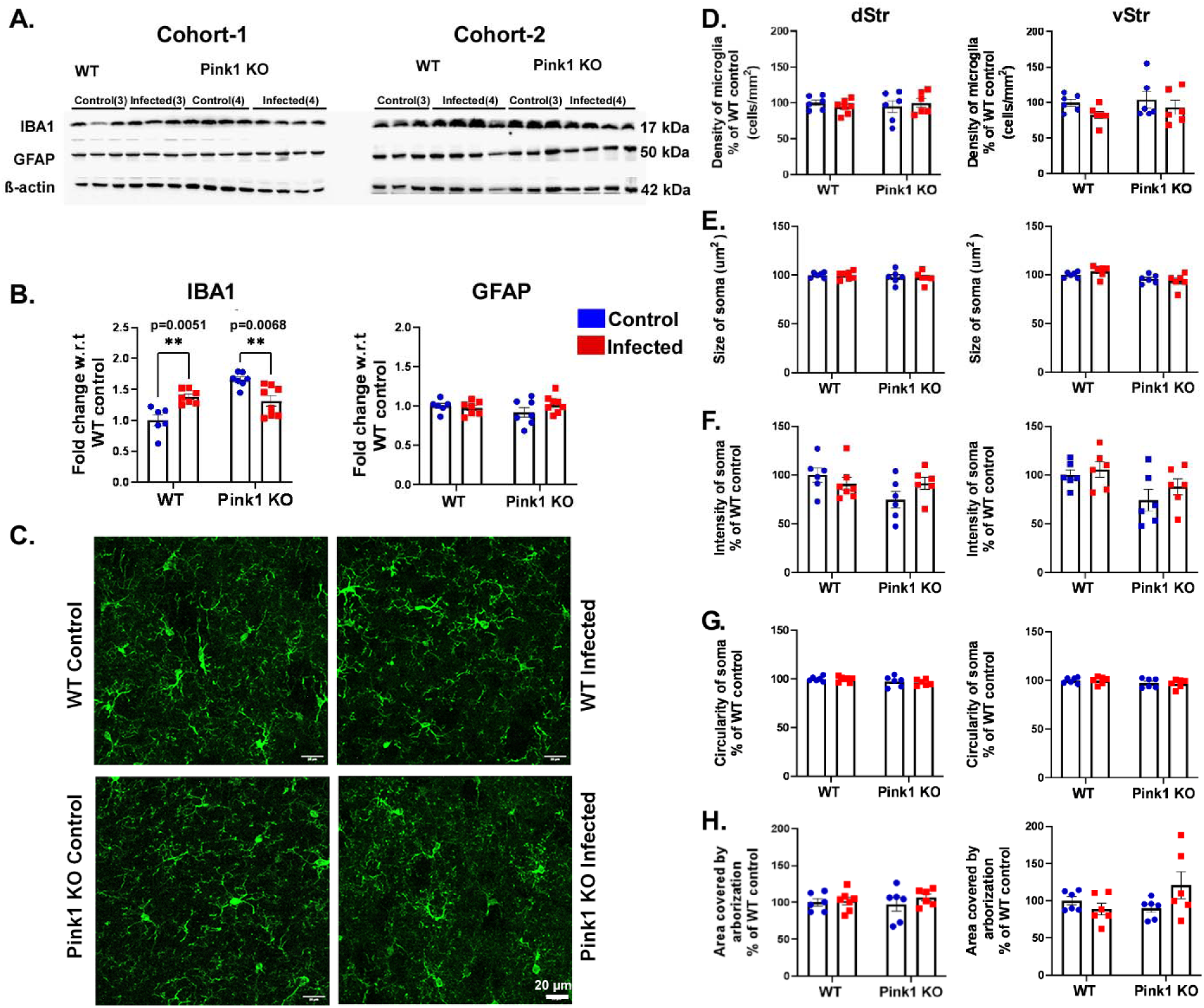
Peripheral infection alters microglial activation marker expression without detectable morphological changes in the striatum at day 13 post infection. Striatal lysates were examined using western blot analysis, targeting IBA1 (microglial marker) and GFAP (astrocytic marker) (A). The blots were quantified and relative expression of IBA1 and GFAP are displayed in B. IBA 1 expression in the striatum was also examined by immunofluorescence staining and quantification. Representative confocal images are shown in (C), scale bar 20μm. Quantification of the density of microglia (D), size of their soma (E), intensity of IBA1 signal (F), circularity (G) and arborization area (H) in both dorsal and ventral striatum was performed. Data represent two independent cohorts of mice, and the results are presented as mean ± S.E.M. Statistical significance was determined using two-way ANOVA with Tukey’s post hoc test at a 95% confidence level.

**Fig 10.**
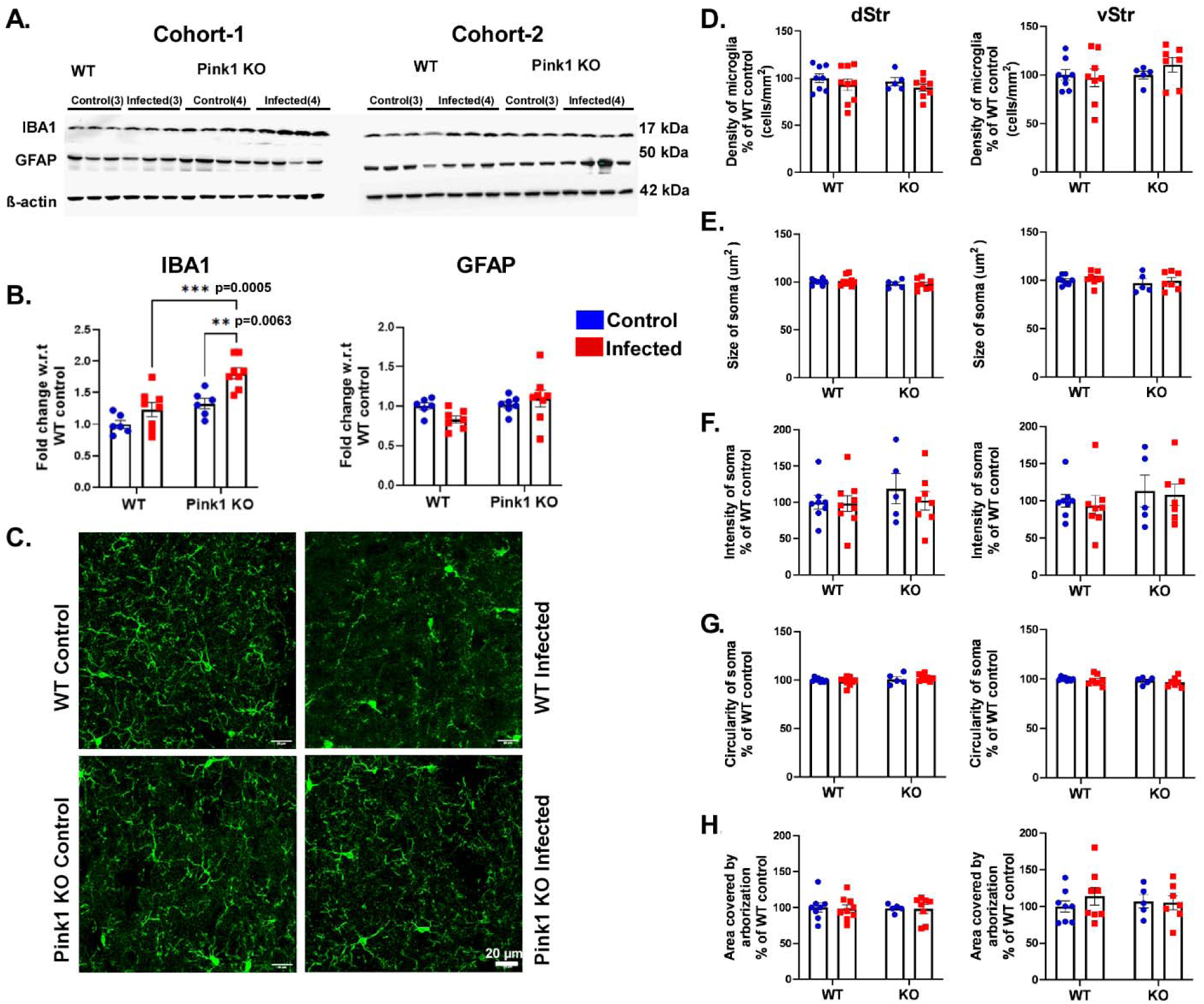
At day 26 post-*Citrobacter rodentium* infection, microglial activation is observed in the striatum, whereas astrocytes remain unaffected. Striatal lysates were analyzed using western blot to detect IBA1 (marker for microglia) and GFAP (marker for astrocytes) (A). The blots were quantified, and the relative expression levels of IBA1 and GFAP are shown in (B). Additionally, IBA1 expression in the striatum was assessed through immunofluorescence staining and quantification. Representative images are presented in (C) scale bar 20μm. Quantitative analysis included microglial density (D), soma size (E), IBA 1 signal intensity (F), circularity (G), and arborization area (H) in both the dorsal and ventral striatum. Data are derived from two independent cohorts of mice and are the results presented as mean ± S.E.M. Statistical significance was determined by two-way ANOVA with Tukey’s post hoc test at a 95% confidence level.

In a separate immunohistochemical experiment, IBA1 and CD68 co-immunostaining was performed to further assess whether activated microglia were engaged in phagocytic activity. The number of IBA1+ cells, as well as CD68 intensity and density within IBA1+ microglia, were quantified in striatal sections. No significant differences were observed in either IBA1+ cell counts or CD68 staining patterns between WT and KO infected mice compared with controls (S2 Fig). The absence of alterations in IBA1+ microglial density and CD68 expression suggests that the infection does not promote phagocytic activation of microglia in the striatum, indicating limited involvement of innate immune mechanisms under these conditions.

### Immune cell profiling of brain and spleen reveals neutrophil proliferation and cerebral infiltration in both WT and Pink1 KO mice after *Citrobacter* infection

Our findings of spleen enlargement, BBB permeability increases, and systemic inflammation accompanied by elevation of the levels of cytokines and chemokines (CXCL-1, IP-10 and IL-17) known to promote the recruitment of peripheral immune cells suggest the possibility that *Citrobacter* infection might promote entry of peripheral immune cells into the brain. To examine this, we characterized the immune cell subpopulations in the spleen that proliferated upon intestinal infection and evaluated whether this was accompanied by altered infiltration of these cells into the brain at different time points. Flow cytometry was used to examine the presence of CD4+ T cells, CD8+T cells, NK cells, B cells, eosinophils, neutrophils, inflammatory monocytes, macrophages and dendritic cells in the spleen and in the brain. The gating strategies used are shown in S3 and S4 Figs.

In the spleen at day 13 post infection, neutrophil populations were elevated in the infected mice, irrespective of genotype. A similar trend persisted at day 26, although this effect decreased and was non-significant (Fig 11 A). At day 26 post infection, we observed a significant increase in the abundance of B cells and a decrease of the abundance of CD8+ T cells in the spleen of both Pink1 WT and KO infected mice (Fig11B).

**Fig 11.**
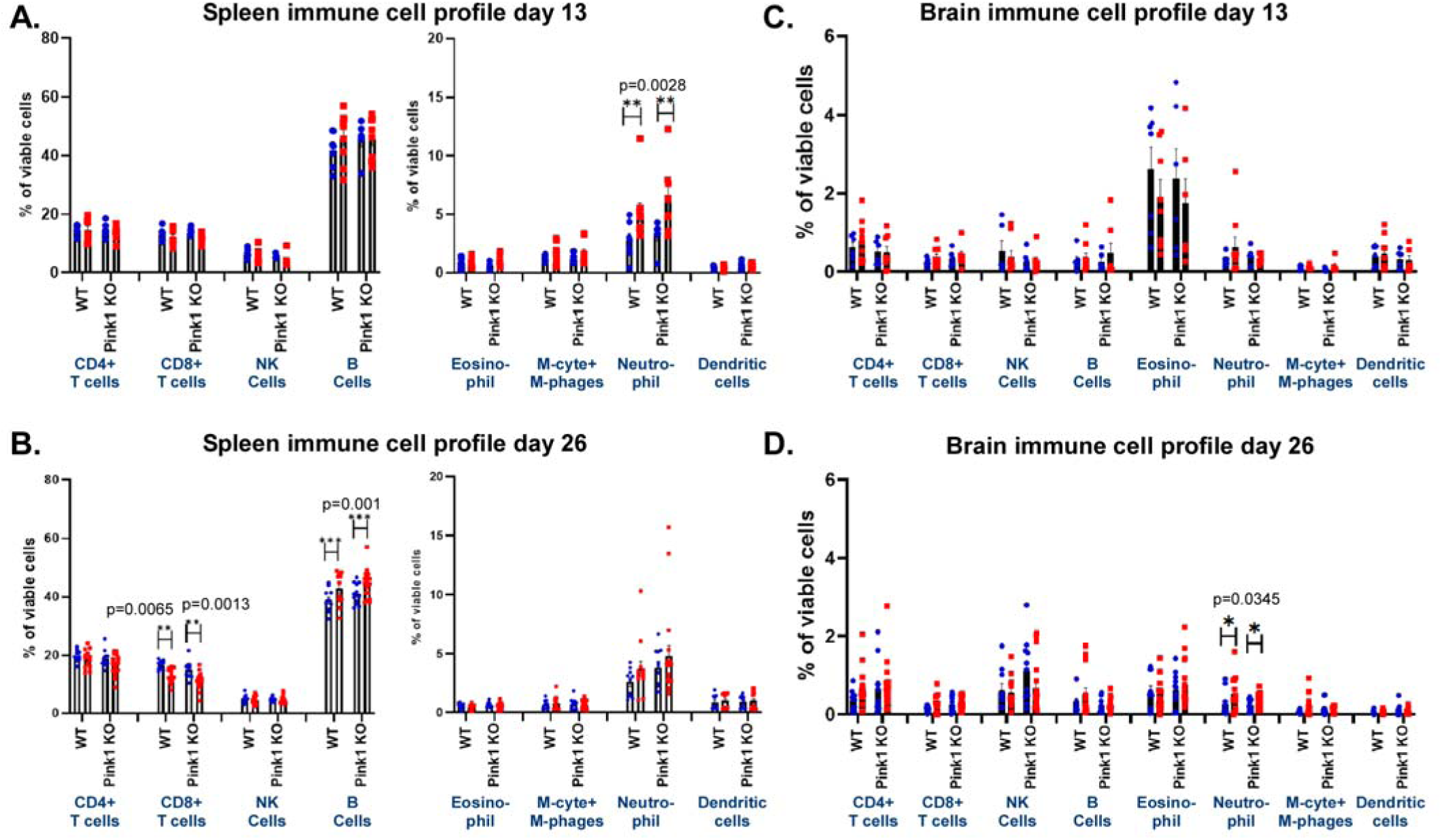
Immune cell profiling of the spleen and brain shows similar increases in neutrophils in the periphery at day 13 and cerebral infiltration of neutrophils at day 26 post-infection in both wild-type (WT) and PINK1 knockout (KO) mice following *Citrobacter* infection. Spleen and brain lysates from all experimental groups were stained with a viability dye and fluorophore-conjugated antibodies to identify CD4+ T cells, CD8+ T cells, NK cells, B cells, eosinophils, neutrophils, inflammatory monocytes, macrophages, and dendritic cells. The respective immune cell populations were measured by flow cytometry as a percentage of viable immune cells. The immune cell profiles in the spleen at days 13 and 26 post-infection are shown in (A) and (B), respectively, while the profiles in the brain at the same time points are shown in (C) and (D). The name of the immune cells examined for both panel C and D are listed at the bottom of panel D. Data represent three independent experiments, and the results are presented as mean ± S.E.M. Statistical significance was determined using two-way ANOVA with Tukey’s post hoc test at a 95% confidence level. Abbreviations: M-cyte: inflammatory monocytes, M-phages: macrophages.

In the brain, no significant immune cell infiltration was detected on day 13 after the infection (Fig 11C). However, on day 26, a significant elevation of the infiltration of neutrophils was observed in the brain of both Pink1 WT and KO infected mice (Fig 11D). Together these results suggest that a *Citrobacter* infection leads to generation of signals that promote the infiltration of peripheral immune cells in the brain.

### The absence of PINK1, when combined with bacterial infection, increases the activation of inflammatory and immune signaling pathways in the striatum

A final set of experiments were performed to obtain a broader unbiased view on how the brain of WT and Pink1 KO mice responded to the *Citrobacter rodentium* infection. For this, mRNA was extracted from the striatum of the mice at day 26 after the infection and subjected to sequencing. To analyze transcriptome-wide changes amongst the experimental groups, principal component analysis (PCA) was conducted on normalized read counts of expressed genes (read counts > 10). The PCA plots, illustrating the clustering and variance among samples, were generated using the PCA function in Python, providing insights into the underlying differences in transcriptomic profiles of non-infected and infected mice (S5A, S5B Fig). Samples that did not cluster closely with others from the same treatment group in the PCA plot were considered outliers and excluded from the analysis. Volcano scatter plots were generated to compare the DEGs in non-infected and infected WT and KO groups. Differentially expressed genes (DEGs) were identified using DESeq2, with stringent thresholds of p-value < 0.05, log2 fold change > 1, and an average basal expression > 10, ensuring the inclusion of actively expressed genes while filtering out non-expressed ones (S5C, S5D Fig).

The striatal transcriptome profile of Pink1 KO mice was found to be distinct compared to their WT counterparts, highlighting significant biological alterations. Gene Ontology (GO) analysis of differentially expressed genes revealed pathways that are distinctly regulated between WT and KO mice under both non-infected and infected conditions (Fig 12 Ai, ii). Each pathway is organized hierarchically to highlight relationships between processes, with nodes indicating distinct biological functions. Interestingly, the dominant clusters for the infected samples (WT and KO) involve pathways related to synaptic functions and vesicle dynamics, such as “establishment of protein localization to postsynaptic membrane,” “neurotransmitter receptor transport,” “synaptic vesicle membrane organization”, “protein localization to synapse,” and “vesicle-mediated transport” (Fig12 Aii). These processes are critical for neurotransmitter signaling, synaptic plasticity, neuronal communication, protein trafficking and vesicle dynamics. Globally, this finding suggests that these pathways, otherwise known to be perturbed in PD, are affected by the infection. Genes associated with microglial activity (Aif1, CX3CR1, Trem2, P2RY12), were not significantly differentially expressed (Fig12 Bi, ii). Similarly, tight-junction protein-related pathways showed comparable expression between WT and KO mice, both at baseline and during infection. This stability is compatible with the conclusion that blood-brain barrier integrity was not drastically affected and only transiently perturbed to facilitate the entry of peripheral immune cells (Fig 12 Bi, ii).

**Fig 12.**
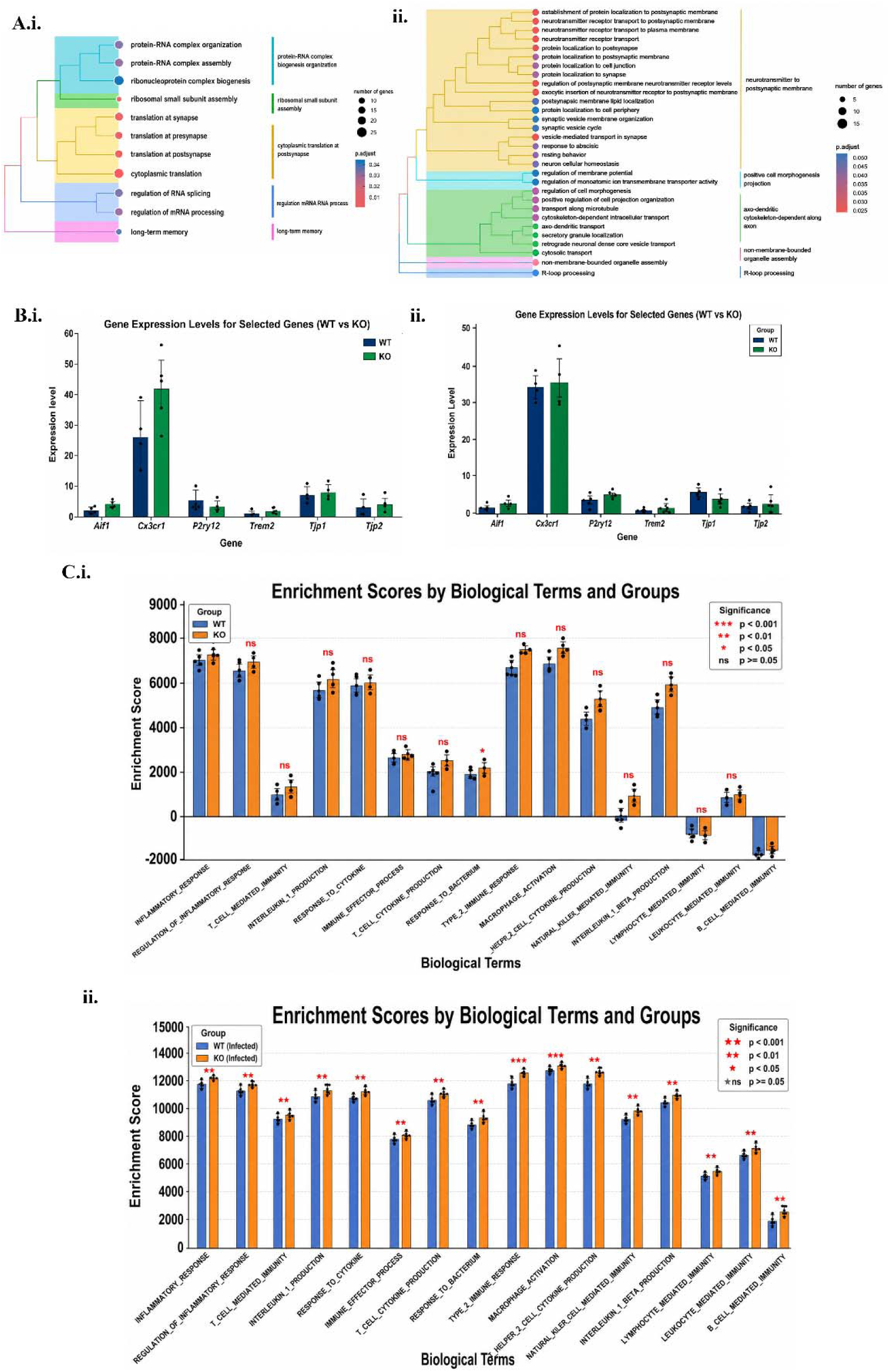
Upregulation of inflammation-related striatal transcription in Pink1 KO mice following *Citrobacter rodentium* infection. The striatal transcriptional profile of uninfected (i) or infected (ii) Pink1 KO and WT mice was compared. Gene Ontology (GO) analysis of differentially expressed genes was examined under both non-infected (Ai) and infected conditions (Aii). Hierarchical clustering grouped enriched biological pathways based on shared gene membership. Each node represents an enriched GO term, with node size proportional to the number of genes associated with the pathway and node color indicating the adjusted P value among the significantly enriched pathways (adjusted P value < 0.05), where lower adjusted P values are shown in red and higher adjusted P values are shown in blue. The clustering of the non-infected (Ai) and infected samples (Aii) identifies different biological pathways differentially expressed by WT and Pink1 KO brain tissue; the findings suggest an impact of the treatment on gene expression, specifically for pathways related to neurotransmitter release and post-synaptic membrane modulation. Microglia-related gene expression remained largely unchanged across conditions (Bi,ii), as did tight junction-related genes. Inflammation-related pathways were significantly upregulated in infected KO mice compared to their WT counterparts, supporting a role for Pink in immune response modulation (Ci, ii). Statistical significance was validated using t-tests on enrichment scores.

To further examine whether the increased BBB permeability observed after *Citrobacter rodentium* infection was associated with transcriptional changes in barrier-associated genes, we interrogated the RNAseq dataset for tight junction markers using normalized read counts prior to differential expression analysis. Expression levels of key tight junction genes, including *Ocln*, *Cldn3*, *Cldn5*, *Cldn12*, *Tjp3* and *Cdh5*, were comparable between infected and non-infected mice across both Pink1 WT and KO genotypes (S6 Fig). Statistical analysis using unpaired t-tests revealed no significant differences for any of these genes. These findings are consistent with the western blot data and suggest that *Citrobacter* infection does not induce overt transcriptional downregulation of major endothelial junction components in the striatum.

Additionally, a closer examination of inflammation-related pathways revealed that although they were not different between WT and Pink1 KO in control animals (Fig 12 Ci), they were significantly increased in the infected KO mice, in keeping with the hypothesis that Pink1 modulates immune responses (Fig 12 Cii). Statistical significance was confirmed by t-tests applied to ssGSEA enrichment scores. These findings, together with our other observations collectively point to altered immune dynamics in Pink1 KO mice after infection, emphasizing their heightened microglial activity and inflammatory predisposition, which could have implications in the long-term for DA neuron function and vulnerability to subsequent age-related or environmental perturbations.

## Discussion

PD, with its broad range of motor and non-motor symptoms is increasingly recognized as a systemic disease involving not only the brain, but also other organs including the gut and the peripheral immune system. It is likely to result from a complex interplay between genetic and environmental factors. Intriguingly, investigations of some of the genetic mutations associated with familial forms of PD are starting to reveal a critical role of multiple PD-related gene products in immune mechanisms. This includes notably Pink1, Parkin, LRRK2 and GBA1 (20–22). In addition, epidemiological studies have highlighted the existence of links between exposure to bacterial or viral pathogens and the risk of developing PD (8, 9, 23–25) . Such results underscore the multifactorial nature of the disease and emphasize the existence of intricate interactions between genetic susceptibility factors and peripheral changes, including in the microbiome, which are thought to act synergistically to affect the onset and progression of PD. Therefore, more work is required to investigate how alterations in intestinal microbiota can contribute to triggering the dysfunction and degeneration of vulnerable neuronal populations in brain, including DA neurons. Following up on recent work showing that gastrointestinal infection with *Citrobacter rodentium* can lead to PD-relevant behavioral and neuroanatomical changes associated with immune cell infiltration in the brain of Pink1 KO mice (13), the present study aimed to obtain a better understanding of the initial effects on the brain of intestinal *Citrobacter rodentium* infection in a Pink1 WT and KO mouse model.

Considering previous work suggesting the existence of a connection between the gut microbiome and BBB integrity (26), we hypothesized that *Citrobacter* gut infection could perhaps lead to an increase in BBB permeability and that in the context of Pink1 loss of function, this could contribute to promoting a state of brain inflammation and elevated peripheral immune cell entry in the brain. Using high resolution, contrast-enhanced pre-clinical MRI to measure BBB permeability, we found that at day 26 after the infection, both Pink1 WT and KO mice show increased BBB permeability in the striatum, thalamus, dentate gyrus and primary somatosensory cortex. This observation is compatible with previous work showing that infection of mice with *Streptococcus agalactiae*, known to cause meningitis, leads to enhanced BBB permeability (27). The same strain has been shown to perturb tight junction proteins in iPSC-derived brain microvascular cells (28).

In the present work, we have not been able to identify the specific mechanism leading to an increase in BBB permeability at day 26 following infection. However, we can exclude that this effect was driven by large-scale alterations in the expression of endothelial tight junction components, as we did not observe robust changes in the protein levels of ZO-1 and ZO-2, two critical scaffolding proteins of the tight junction complex. Consistent with these findings, transcriptomic analysis of striatal RNA-seq data revealed no significant differences in the expression of multiple tight junction genes, including *Ocln*, *Cldn3*, *Cldn5*, *Cldn12*, *Tjp3*, and *Cdh5*. Together, these results suggest that the increased permeability to gadolinium is unlikely to have resulted from pronounced loss or transcriptional downregulation of junctional proteins, but may instead reflect more subtle functional modifications of the endothelial barrier, such as junctional remodeling or inflammatory signaling mediated changes in permeability. Further studies will be required to delineate the precise mechanisms underlying these BBB alterations.

*Citrobacter* gut infection in our model led to peripheral changes in the levels of cytokines and chemokines. Measurements of serum cytokines and chemokines revealed a significant up-regulation of IL-17 and IP-10 at day 13 after the infection, and of IL-17 and CXCL1 at day 26, with CXCL1 showing an elevation only in the infected Pink1 KO mice. Previous evidence shows that inflammatory cytokines and chemokines in the bloodstream can penetrate the brain, contributing to the development of various neuroinflammatory pathology in the context of ischemia, multiple sclerosis, and hypertension (29–31). IL-17A is one of the main cytokines known to disrupt the functioning of the BBB in multiple *in vitro* and *in vivo* rodent models of multiple sclerosis (32, 33). Peripheral inflammatory signals in the infected mice could perhaps directly contribute to the observed changes in BBB permeability. Compatible with this hypothesis, previous work has shown that peripheral cytokines like IL-6, IL-1β, IFN-γ can facilitate BBB permeability in various mouse, rat and zebrafish models (34–36).

Elevations of peripheral cytokines and chemokines have been shown to be associated with microglial activation and DA neuronal loss in the MPTP mouse model (37). Interestingly, in people with PD, the levels of peripheral cytokines and chemokines are positively correlated with the severity of the non-motor symptoms (38). This could perhaps occur because microglial activation leads to alterations in neuronal functions including neurotransmitter release (39). Compatible with a link between cytokine/chemokine elevation and microglial activation, we observed an increase in multiple proinflammatory mediators in the stratum of the infected mice at day 13 after the infections, irrespective of genotype. Such signals decreased close to baseline at day 26, suggesting that the inflammatory signals were short lasting. Such signals could nonetheless contribute to microglial activation that could outlast these initial signals. In line with this, we found that at 26-days post infection, higher IBA1 levels were detected in the striatal lysates in the infected mice. Interestingly, infected Pink1 KO mice were found to have significantly more IBA1 expression compared to the WT infected mice. Also, we observed a tendency of higher IBA1 expression in the non-infected Pink1 KO mice compared to the WT non-infected, which might reflect an existing higher number of microglia in the brain of the Pink1 KO mice; this could lead to the establishment of a chronic state of microglial activation in the brain of the Pink1 KO mice upon bacterial infection. Microglial activation has been proposed to be a critical contributor to DA neuron degeneration (40–42). Interestingly, combined biochemical and histological analyses in our study provide important clarification regarding the nature of the myeloid response observed following infection. While western blotting of striatal lysates revealed a modest increase in total IBA1 levels, region-specific immunofluorescence analyses demonstrated no significant changes in microglial density, morphology, or CD68 expression within IBA1+ cells. Since CD68 is widely used as a marker of lysosomal and phagocytic activation associated with reactive microglia (43–45), the absence of altered CD68 intensity or distribution suggests that resident parenchymal microglia do not undergo robust functional activation. Importantly, bulk tissue lysates might capture signals from multiple cell types, including perivascular macrophages and infiltrating monocytes present in the striatum. Therefore, the increase in IBA1 observed by western blot likely reflects contributions from these non-parenchymal populations along with resident microglia within the striatal tissue. Together, these results highlight the fact that infection induces only mild myeloid changes without promoting phagocytic microglial activation in the striatum.

Among the mediators we found elevated at day 13 after infection in the striatum is IFN-γ, an inflammatory cytokine that has been reported to promote BBB damage in viral encephalitis and meningitis models (46–48). Additionally, IFN-γ is a well-known activator of microglia (49).

Therefore, induction of IFN-γ could perhaps trigger a cascade leading to subsequent microglial activation at later time points. Further work will be required to test this. Although we did not identify the origin of the cytokines and chemokines found to be elevated in the striatum after the infection, it is possible that some of these are secreted by endothelial cells. Previous work has shown that brain endothelial cells can transport and secrete cytokines (50, 51). Therefore, we can postulate the implication of a sequence of events linking peripheral cytokines and chemokines elevation to subsequent limited microglial activation and release of other signals derived from endothelial cells and/or pericytes. Further experimentation will be needed to establish the cause-and-effect relationships predicted by this model.

Some of the cytokines and chemokines elevated in the brain of the Pink1 WT and KO mice could act to promote brain entry of peripheral immune cells. Such entry of peripheral immune cells, including CD8+ T cells, has been suggested to be critical to cause the degeneration of DA neurons in Pink1 KO mice after repeated *Citrobacter* infections (13, 14). Here we therefore examined whether a single infection with *Citrobacter* was sufficient to promote immune cell entry in the brain. Using flow cytometry-based profiling of spleen and brain immune cells, we found an amplification of neutrophils in the spleen of infected mice at day 13 post infection, and increased levels of neutrophils in the brain at day 26 post infection. Interestingly, a higher neutrophil to lymphocyte ratios has been identified as a potential biomarker for PD (52). In line with such results, we observed an increase in neutrophils and a decrease in CD8+ T cells in the spleen post infection. Although the significance of a drop in peripheral T cells in PD is still a matter of debate (53, 54), mounting evidence suggests that in the brain, an activation of microglia and the entry of CD4 or CD8 T cells acts as an initiating factor for neurodegeneration (14, 55–60). While in our study, flow cytometry indicated the presence of neutrophils within brain homogenates, histological validation using immunostaining would provide important spatial confirmation of neutrophil infiltration and regional localization. However, such analyses were not performed in the present study, representing a limitation of our work. Future studies incorporating immune cell imaging in brain sections will be necessary to more definitively assess immune cell entry into the brain following infection.

It can be considered unlikely that a single gastrointestinal infection with a relatively mild pathogen such as *Citrobacter rodentium* (analogous to a self-limiting gastro-intestinal infection in humans) would result in marked neurodegeneration. In line with this, our western blot and immunohistochemical investigation of markers of the DA system in the striatum did not reveal any significant decreases after the infection. We also did not detect any significant decreases in such markers when comparing non-infected WT and Pink1 KO mice, consistent with previous reports of a lack of DA system degeneration in Pink1 KO mice (61, 62). Additionally, quantitative analysis of tdTomato-positive DA neurons in both the SNc and VTA revealed no significant reductions in neuronal numbers in either WT or Pink1 KO mice at day 26 post-infection. These findings indicate that the observed vascular and inflammatory alterations occur in the absence of dopaminergic neurodegeneration. However, positron emission tomography (PET) and single-photon emission computed tomography (SPECT) imaging of early-onset PD patients with Pink1 mutations has shown presynaptic DA system dysfunction in the striatum, similar to that seen in late-onset idiopathic PD patients (63, 64). Together, these results support the hypothesis that inflammation deriving from infection in the present model induced early cerebrovascular perturbations and neuroimmune activation without measurable neurodegeneration. This is consistent with accumulating evidence in PD suggesting that BBB disruption and neuroinflammation are early pathological events that can contribute to neuronal vulnerability over time. The absence of DA neuron loss at this stage suggests that the pathological changes observed here may reflect a prodromal-like phase, during which vascular and immune mechanisms prime the neural environment for subsequent neurodegenerative processes. We also hypothesize that repeated cycles of inflammation and immune cell infiltration into the brain may progressively create neurotoxic conditions that contribute to DA neuron damage.

Loss of function of Pink1 has been proposed to disinhibit mitochondrial antigen presentation and to enhance innate immune mechanisms (65, 66). In the present work, although we found equivalent changes in BBB permeability at day 26 in Pink1 WT and KO mice, we did obtain evidence for a larger impact of the infection on inflammatory mechanisms in the KO mice. We found that the colon index at day 26 after the infection was still significantly elevated in the Pink1 KO mice, while it had recovered in the WT mice, suggesting a long-lasting inflammatory state in the KO mice. We also found a persistent elevation of IL-17 in the serum of the Pink1 KO mice at day 26 after the infection, here again with a faster recovery in the WT mice, and a significant elevation of KC (CXCL1) at day 26 after the infection only in the KO mice. Finally, we found a significant elevation of IL-17 in the striatum of the Pink1 KO mice but not in WT mice at day 13 after the infection. Together these observations make it plausible that over time, with repeated cycles of inflammation, mice or people with Pink1 loss of function may suffer more important consequences, in line with our previous observation of increased loss of DA system innervation in the striatum of Pink1 KO mice exposed to 4 cycles of *Citrobacter rodentium* infection (13). Our analysis of striatal transcriptome, revealing elevated enrichment scores for biological pathways related to inflammation and immune regulation, is also in keeping with this conclusion. These results suggest a heightened inflammatory state in the brain of infected KO mice compared to infected WT. Over time, increased microglial activation and inflammation may contribute to neuronal dysfunction and damage, compromising neuronal health.

Together, our data indicate that *Citrobacter* infection can trigger inflammatory responses both in the periphery and the CNS, which perturbs BBB integrity, facilitates the entry of immune cells into the CNS and leads to brain inflammation, potentially initiating or exacerbating pathophysiological processes leading to neurodegeneration in PD (Fig 13). Collectively, our results are compatible with the possibility that PD-related pathophysiological mechanisms derive from an intricate relationship between genetic vulnerability factors and peripheral mechanisms, including the state of intestinal tissues and the functional integrity of the BBB, together influencing brain inflammation, including the state of microglial cells and the entry of peripheral cells in the brain. The present work highlights the need for further exploration of the implication of the gut-brain axis in PD pathogenesis and progression, holding the promise for developing innovative therapies for improving the quality of life of people with PD.

**Fig 13.**
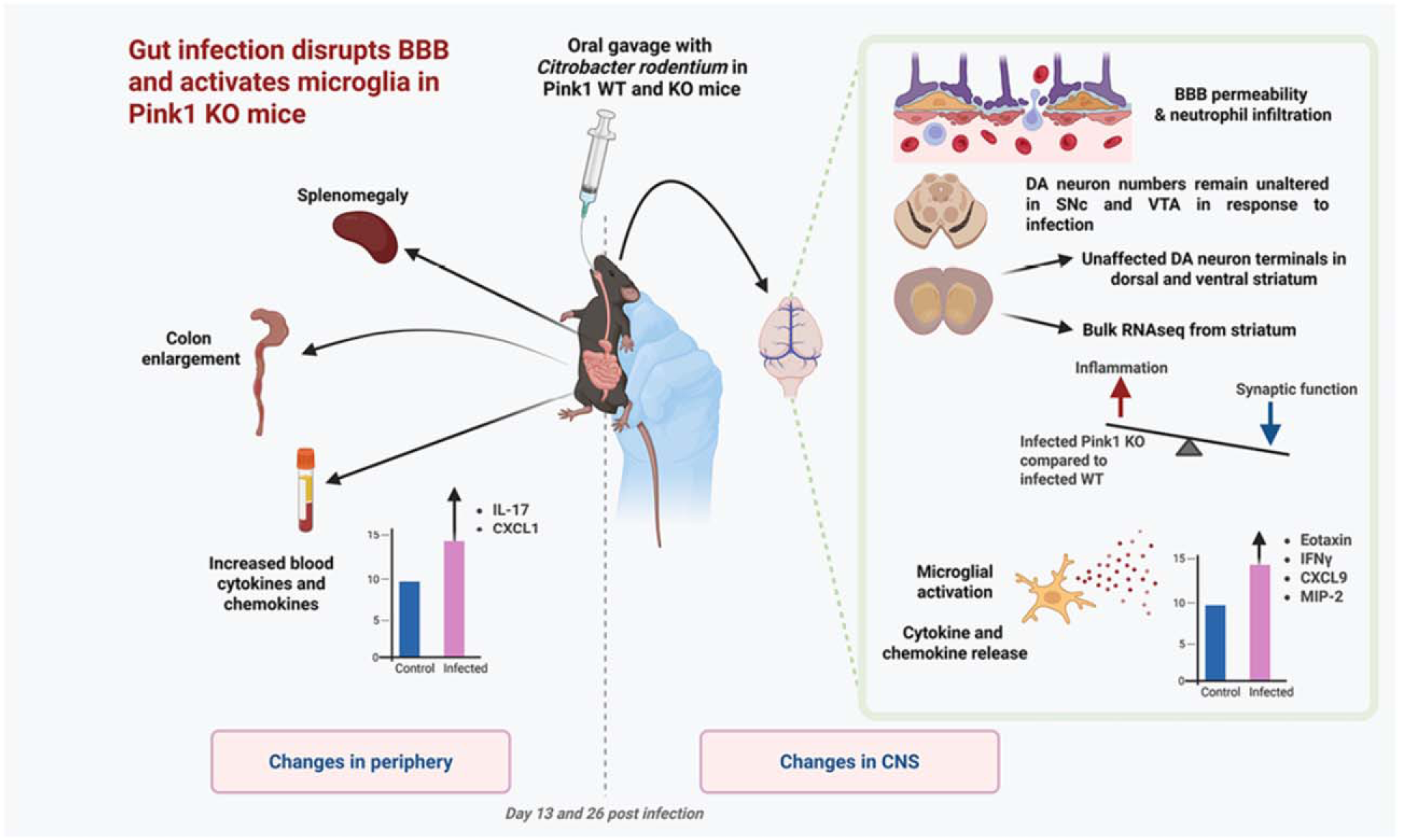
Summary of systemic and central effects of Citrobacter rodentium infection in Pink1WT and KO mice. Oral gavage with *Citrobacter rodentium* induces peripheral immune activation characterized by splenomegaly, colon enlargement, and increased circulating cytokines and chemokines. In the central nervous system, infection is associated with increased BBB permeability as detected by MRI and elevated neutrophil signals in the striatum. Bulk RNA-seq analysis of striatal tissue reveals upregulation of inflammation in the KO mice. Quantification of dopaminergic neurons in the substantia nigra pars compacta (SNc) and ventral tegmental area (VTA), as well as dopaminergic terminals in the striatum, shows preservation of neuronal integrity following infection. Together, these findings indicate that a single peripheral infection induces modest neurovascular and neuroimmune alterations without overt dopaminergic neurodegeneration. Created in BioRender. Mukherjee, S. (2026). https://app.biorender.com/citation/69821547900d887bb43544d4.

In closing, an important limitation of the present study is that analyses were restricted to a single infection paradigm. Previous work (67) has suggested that repeated inflammatory challenges with *Citrobacter* can produce more pronounced and persistent neuroimmune alterations than a single transient exposure. Thus, while we observed only modest myeloid changes following one episode of infection, it remains possible that multiple or recurrent infections could elicit stronger microglial activation, enhanced BBB dysfunction, or more substantial neurodegeneration, particularly in the context of Pink1 deficiency. Future studies directly comparing single versus repeated infection paradigms across cellular, molecular, imaging, and immune readouts will be essential to distinguish transient responses from cumulative pathological effects and to better model chronic inflammatory “second hit” scenarios relevant to PD.

## Material and Methods

### Animal breeding

Mice were housed and maintained according to protocols approved by the animal ethics committees of the Université de Montréal (CDEA) and McGill University. Guidelines defined by Canadian Council of Animal Care were strictly followed for all experiments. Mice were maintained in a 12h light/dark cycle with defined temperature (21 °C) and humidity (60%) and adequate access to food and water.

Pink1 KO mice were a kind gift from Dr. Hansruedi Bueler (RRID:IMSR_JAX:017946). These mice were crossed with DAT-IRES-cre mice (Jackson Laboratory, RRID: IMSR_JAX:006660), driving expression of cre-recombinase under control of the DA transporter (DAT) promoter, allowing selective expression of Cre in DA neurons. Pink1 KO mice were then crossed with Ai9/tdTomato mice (Jackson Laboratory, RRID: IMSR_JAX:007909) in which a loxP-flanked STOP cassette is designed to allow transcription of tdTomato in cells expressing cre recombinase. Pink1 HET;DAT-IRES-Cre mice were then crossed with Pink1 HET;Ai9/tdTomato mice to generate offsprings expressing tdTomato selectively in DA neurons within the Pink1-deficient background. The final mice generated were either KO or WT for Pink-1, and carried one allele of cre and one allele of tdTomato, thereby enabling selective expression of the tdTomato fluorescent protein in DA neurons. These mice are thereafter called DATcre;Ai9;Pink1 or for short, DATcrePink1 mice. The genotyping of these mice was performed with a KAPA2G Fast Hot Start DNA polymerase kit (Roche, Cat# KK7352). The following primers were used to identify Pink1 WT and KO mice: TCCCTCTATGGCGTCCTCTT (WT, forward), GCAACTGCAAGGTCATCA (WT, reverse), GCACCCTGACCTTGGTTCCTA (KO, forward), GGGGGAACTTCCTGACTAGG (KO, reverse) (Integrated DNA Technologies). Both male and female mice were used in the present study. The results obtained from both sexes were pooled because the study was underpowered to assess sex-dependent differences.

### *Citrobacter* infection in DATcrePink1 mice

The enteric bacteria *Citrobacter rodentium* (ATCC, Cat #51459), was cultured and maintained under biosafety level 2 equipment and facilities at McGill University. The chloramphenicol resistant strain DBS 100 was used for infecting mice. Briefly, on day 1, the bacteria were taken from a frozen stock, and a streak-plate was prepared on LB agar supplemented with chloramphenicol and left overnight at 37 °C. On day 2, a single colony was picked and inoculated in 3 ml of LB-broth and allowed to grow for 17.5h at 220 RPM at 37 °C. On the day of infection, 200µl from the overnight culture was used for oral gavage for each mouse using a 22G gavage needle. Similarly, the control mice received an equal volume of LB without any bacteria. At the same time, serial dilution of the culture was made, and plated on LB-agar. The next day, colony forming units (CFU) were monitored along with quantification of the actual doses of bacteria given to the mice. Routinely, in all experiments, the CFU of gavage in mice was found to be within the range of 10^8^-10^9^.

At day 6 post gavage, fecal pellets were collected from each of the infected mice, weighed and homogenized in 1 ml PBS using a MagnaLyser at speed 6000 RPM for 1 min. Serial dilutions were prepared and undiluted to 10^-9^ dilutions were plated on MacConkey agar plates with chloramphenicol to detect the bacterial load. After overnight incubation at 37 °C, colony counting was performed from the dilution plates and CFU/g of feces was calculated for each infected mouse. No mortality or severe adverse events were observed in the present study.

### Magnetic Resonance Imaging (MRI) of the brain and image processing

All procedures were carried out according to animal health and safety protocols of McGill University. Imaging was performed using the Bruker Pharmascan preclinical 7 Tesla MRI scanner (Billerica, MA, USA) of the McConnell Brain Imaging Centre (Montreal, Quebec). Briefly, anaesthesia in mice was induced and maintained with isoflurane (2-3%) during MRI data acquisition. A relaxation enhancement at variable repetition time (RARE-VTR) acquisition protocol for T1 mapping was implemented with the following parameters: 150×150×500 μm^3^; field of view; 133×133×18 matrix size; TR = [8000, 3600, 2400, 1480, 940, 650, 501.1] ms. An anisotropic acquisition was selected to maximize in-plane resolution, given the time constraints related to maintenance of animal anaesthesia. The scan was repeated twice: once prior to and once following gadolinium-based contrast agent (GBCA) injection via the tail vein with a 0.4 mmol/kg dose in each mouse. The GBCA was diluted in saline. Successful delivery of GBCA into systemic circulation of each animal was verified by acquiring a single RARE acquisition pre- and post-injection.

Reconstructed image sets were motion-corrected by computing a rigid (6 degrees of freedom) transformation between the TR = 8000 ms and all subsequent volumes. Images were denoised using the Marcenko-Pastur principal component analysis (MP-PCA) (68) and corrected for Gibb’s ringing (69) using algorithms implemented in the DiPy package version 1.4.0 (Python 3.8, RRID:SCR_008394) (https://github.com/dipy/dipy) (RRID:SCR_000029) (70). The brain was segmented from the skull and surrounding soft tissue using the Shape descriptor selected extremal regions after morphologically filtering (SHERM) algorithm implemented in MATLAB (version R2019b, http://www.mathworks.com/products/matlab/, RRID:SCR_001622) (https://github.com/liu-yikang/SHERM-rodentSkullStrip/tree/master) (71). Following brain extraction, bias field correction was applied to the brain portion of the TR = 8000 ms image. The estimated bias field was then used to correct all subsequent volumes. Bias field estimation was carried out using the N3 algorithm implemented in ANTs software available at https://scicomp.ethz.ch/public/manual/ants/2.x/ants2.pdf. The extracted and corrected mouse brain image was then registered to a high resolution (40×40×40 μm^3^) anatomical template (72) using affine registration (12 degrees of freedom). The resultant transformation matrix was inverted and used to register template labels to the native space of the RARE-VTR acquisition for region of interest (ROI) parcellation. All image-space transformations were computed using ANTs. Conventional voxel-wise fitting of the mono-exponential saturation recovery curve sampled by the RARE-VTR protocol was carried out using a nonlinear least squares solver (lsqnonlin) implemented in MATLAB vR2019b. This fitting procedure was used to produce whole brain T1 maps pre- and post-GBCA injection. Putative GBCA-induced T1-shortening was evaluated by constructing histograms of T1 values within each ROI and computing the earth mover’s distance (EMD) (73) between corresponding histograms.

### Immunohistochemistry in brain slices

At days 13 and 26 post infection, *Citrobacter*-infected and non-infected mice were anaesthetized with pentobarbital followed by trans-cardiac perfusion with 1X cold PBS and 4% PFA. 24h after, brain samples were transferred to a 30% sucrose solution for 48h. 40µm serial coronal sections were cut from each brain using a Leica CM1800 cryostat and collected in anti-freeze solution. The floating slices were washed with 1X PBS to remove the anti-freeze solution (3 X 10 min). This was followed by permeabilization and blocking for 60 min under agitation in a solution with 0.3% Triton X, 5% goat serum and 10% BSA. Slices were incubated with the following primary antibodies overnight with agitation: TH (1:1000, Millipore Cat# AB152, RRID: AB_390204), DAT (1:1000, Millipore Cat# MAB369, RRID: AB_2190413), IBA1 (1:1000, FUJIFILM Wako Pure Chemical Corporation Cat# 019-19741, RRID: AB_839504). The next day, slices were washed with PBS followed by appropriate Alexa Fluor 488/647 conjugated secondary antibody (rabbit/rat) [Goat anti-Rabbit IgG (H+L) Cross-Adsorbed Secondary Antibody, Alexa Fluor 488, Cat # A-11008, RRID: AB_143165; Goat anti-Rat IgG (H+L) Cross-Adsorbed Secondary Antibody, Alexa Fluor 647, Cat # A-21247, RRID: AB_141778; both from Invitrogen for 2h at RT with agitation. Following 3 X 10 min wash with 1X PBS, slices were mounted on charged microscope slides (Surgipath X-tra, Leica, 3800200) and stored at 4 °C until imaging.

For the assessment of microglial morphology in relation to CD68 expression, striatal sections were immunostained with primary antibodies against CD68 (1:100, BioLegend, #137001, RRID: AB_2044003), IBA1 (1:1000, Synaptic Systems, #234308, RRID: AB_2924932), and tyrosine hydroxylase (TH) (1:1000, Millipore Cat# AB152, RRID: AB_390204) following the same staining protocol described above. Corresponding species-specific secondary antibodies conjugated to Alexa Fluor 488, 546, and 647 were used for fluorescence detection [Goat anti-Guinea Pig IgG (H+L) Highly Cross-Adsorbed Secondary Antibody, Alexa Fluor 488, Cat# A-11073, RRID: AB_2534117; Goat anti-Rabbit IgG (H+L) Cross-Adsorbed Secondary Antibody, Alexa Fluor 546, Cat # A-11010, RRID: AB_2534077; Goat anti-Rat IgG (H+L) Cross-Adsorbed Secondary Antibody, Alexa Fluor 647, Cat # A-21247, RRID: AB_141778; from Invitrogen].

In addition, series of 40 µm thick mesencephalic sections were immunostained with primary antibodies against TH and red fluorescent protein (RFP) (1:1000, Synaptic Systems, #409006 RRID: AB_2725776) to visualize tdTomato-positive DA neurons. One out of three sections across the series of sections encompassing the SNc and VTA were selected, typically providing six to nine sections per mouse. Sections were subsequently incubated with appropriate secondary antibodies [Goat Anti-Chicken IgY (H+L), Highly Cross-Adsorbed Secondary Antibody CF543, Cat # 20311 from Biotium; Goat anti-Rabbit IgG (H+L) Cross-Adsorbed Secondary Antibody, Alexa Fluor 647, Cat # A-21244 from Invitrogen, RRID: AB_2535812].

### Confocal imaging and image processing

Confocal imaging was performed using an Olympus Fluoview FV1000 point-scanning confocal microscope (Olympus Canada) (RRID:SCR_014215). To minimize nonspecific bleed-through signal, images were acquired sequentially with laser excitation at specific wavelengths: 488 nm, 543 nm, and 633 nm. Image acquisition utilized a 60X, NA 1.42 objective to ensure optimal resolution. In this study, three specific brain slices were selected for analysis. The first slice was chosen at a rostro-caudal location where the corpus callosum from both hemispheres was nearly touching. The second slice was selected at a rostro-caudal location when the anterior commissure formed a straight line, and the third slice corresponded to at a rostro-caudal location where the hippocampus was first seen. Regions of interest within the dorsal and ventral striatum were defined for further analysis. All acquired images were processed using ImageJ software (https://imagej.net/) (RRID:SCR_003070) (74). To facilitate image analysis, custom scripts were developed in Image J (https://github.com/Louis-EricTrudeau/Trudeau-lab/tree/main/Mukherjee-2026). These custom scripts enabled precise and efficient analysis of the acquired images. Prior to quantification, image preprocessing techniques were applied to enhance the accuracy and reliability of the analysis. Two commonly utilized preprocessing methods were employed: “convoluted background subtraction” and “rolling ball background subtraction”. The analysis of TH and DAT involved quantifying fluorescence intensity and surface occupied by signal to provide insights into the expression levels and spatial distribution of these two proteins in the axonal domain of DA neurons.

For the reconstruction and analysis of microglia, z-stack images were captured at intervals of 2 microns. The first step involved segmenting the images to identify cell bodies. This segmentation process was developed in ImageJ, also allowing to trace the cells’ processes. Parameters such as microglia density, soma area, soma fluorescence intensity, soma perimeter, soma circularity and the area and intensity of the processes were quantified.

For Iba1 and CD68 quantification, whole-slide images were acquired using a Zeiss slide scanner (ZEISS Axioscan 7 Digital Slide Scanner, RRID:SCR_027284) with laser excitation at 405, 488, 546, and 647 nm. Three coronal striatal sections per brain were selected as previously described. Each section was imaged across multiple z-planes using a 40X objective with a z-step of 1 µm. Acquired image stacks were deconvolved using the automatic function in Batch Deconvolution software (version 6.10, Nikon) (https://deconv.laboratory-imaging.com/process), to improve signal quality and reduce background noise. Deconvolved images were subsequently processed and analyzed using Nikon NIS-Elements software (version 6.20.02) https://www.microscope.healthcare.nikon.com/products/software/nis-elements/software-resources. An extended depth-of-field (EDF) reconstruction was generated from each z-stack, followed by application of local contrast enhancement and intensity equalization to homogenize signal across samples. To restrict analyses to the striatal region, TH-positive areas were delineated using intensity thresholding based on representative test images. Within these regions, microglial cell bodies were detected and counted using a machine learning–based segmentation approach implemented in NIS-Elements. IBA-positive and CD68-positive objects were further identified using optimized intensity thresholds, and the total area and mean fluorescence intensity for each marker were quantified.

For quantification of mesencephalic DA neurons, whole-slide images were acquired in the same slide scanner with laser excitation at 546 and 647 nm for TH and RFP signals. Coronal midbrain sections spanning bregma −2.92 to −3.88 mm were selected for analysis. Sections were imaged through multiple z-stacks using a 20X objective with a z-step of 1 µm. Image stacks were processed in Nikon NIS-Elements software by generating EDF reconstructions, followed by automated detection of tdTomato (RFP) positive DA neurons using a machine learning-based algorithm. Regions of interest corresponding to the SNpc and VTA were manually delineated. The number of DA neurons for each brain slice was plotted in relation to the corresponding rostro-caudal bregma coordinates, and the total number of neurons within each region was approximated using the area under the obtained curve.

### Western-Blot

The striatum was micro-dissected from the brain of *Citrobacter* infected and non-infected mice at days 13 and 26 post infection, and homogenized (Ultra-Turrax T8, Ika-Werke, Germany) in RIPA buffer (Fisher Scientific, PI89900) containing a protease inhibitor cocktail (P840, Sigma). Homogenized tissue samples were centrifuged at 12000g for 30 min at 4 . Supernatant was collected, and protein quantification was performed with BCA reagent (Thermo Scientific Pierce BCA Protein Assay Kit, PI23227). 20μg of each sample was separated on 8-12% SDS-PAGE followed by transfer onto a nitrocellulose membrane. Membrane blocking was performed with Every Blot blocking buffer (BioRad, Cat #12010020) at RT with gentle shaking for 5 min. The membranes were incubated overnight at 4°C with respective antibodies (Table 1) with gentle shaking. Membranes were washed 5 times with TBST buffer for 5 min each time. After this, appropriate secondary antibodies were added (Peroxidase-conjugated AffiniPure Donkey Anti-Rabbit IgG, 711-035-152/ Peroxidase-AffiniPure Goat Anti-Mouse IgG,115-035-146/ Peroxidase-AffiniPure Goat Anti-Rat IgG, 112-035-003, all three from Jackson ImmunoResearch Laboratories) and the incubation was performed at RT for 90 min with gentle shaking. Membranes were washed again with TBST buffer (5X5 min) and developed using the Clarity Western ECL substrate (Bio-Rad, product #1705061). Images were captured on a luminescent image analyzer (ChemiDoc MP imaging system, BioRad, RRID:SCR_019037). Membranes were stripped and re-probed for β-actin as a loading control. Densitometry analysis was performed in Image J software and the intensities of the genes of interest were normalized to the respective β-actin band intensity of that sample.

**Table 1.** List of primary and secondary antibodies used in Western-Blot experiments.

| Name of antibody | Species | Dilution | Manufacturer | Catalog Number/RRID |
| --- | --- | --- | --- | --- |
| ZO-1 | Rabbit | 1:1000 | Cell signaling technology | Cat# 13663, RRID: AB_2798287 |
| ZO-2 | Rabbit | 1:1000 | Cell signaling technology | Cat# 2847, RRID: AB_2203575 |
| GFAP | Rabbit | 1:2000 | Agilent Technologies Singapore | Z0334, RRID: AB_10013382 |
| IBA1 | Rabbit | 1:1000 | Fujifilm Wako Pure Chemical Corporation | Cat# 016-20001, RRID: AB_839506 |
| Beta actin peroxidase | Mouse | 1:15000 | Sigma | Cat# A3854, RRID: AB_262011 |
| DAT | Rat | 1:1000 | Millipore | Cat# MAB369, RRID: AB_2190413 |
| TH | Mouse | 1:1000 | Millipore | Cat# MAB318, RRID: AB_2201528 |
| Peroxidase-conjugated AffiniPure Donkey Anti-Rabbit IgG | Donkey | 1:5000 | Jackson ImmunoResearch Laboratories | 711-035-152<br>RRID: AB_10015282 |
| Peroxidase-AffiniPure Goat Anti-Mouse IgG | Goat | 1:5000 | Jackson ImmunoResearch Laboratories | 115-035-146<br>RRID: AB_2307392 |
| Peroxidase-AffiniPure Goat Anti-Rat IgG | Goat | 1:5000 | Jackson ImmunoResearch Laboratories | 112-035-003<br>RRID: AB_2338128 |

### Immune cell isolation from brain and spleen

At days 13 and 26 post infection, *Citrobacter* infected and non-infected mice were anaesthetized with pentobarbital. To distinguish the circulating immune cell population from the infiltrating and resident immune cells within the brain, mice were intravenously injected with 3µg of BV711 conjugated anti CD45 antibody (BioLegend Cat# 103147, RRID: AB_2564383). Three min post injection, cervical dislocation was performed, the brain was quickly collected, minced with a scalpel and digested to single cells using 2mg/ml collagenase D (11088882001, Sigma) and 28U/ml DNase I (4536282001, Sigma) for 45 min at 37 . At every 15 min interval, the tissue was triturated using a glass-pipette and a fresh dissociation solution was added. Post digestion, the enzyme activity was neutralized by a buffer containing PBS with 2 mM EDTA and 2% FBS. The dissociated tissue was passed through a 100µm cell strainer (Falcon, 352360) to remove any undigested pieces. The homogenate was centrifuged at RT for 2 min at 200g and 3 min at 300g and the supernatant was carefully discarded. To remove myelin, the pellet was resuspended in 10 ml of 37% (v/v) Percoll diluted in HBSS (gradient density separation solution) and centrifuged at RT for 10 min at 500g without brakes. The layer of myelin was suctioned off and the cell pellet was washed with PBS-EDTA-FBS solution. After this step, cell counting was performed using a hemocytometer followed by staining and analysis in flow cytometer (BD LSRFortessa X-20, RRID:SCR_025285).

The spleen was collected from both infected and non-infected mice at the same time as the brain and kept in ice cold 1X PBS. The tissue was dissociated with the thumb-rest of the plunger of a syringe on a 100µm cell strainer (Falcon, 352360). To harvest the maximum number of cells, the filter was washed several times with dissociation buffer containing PBS-EDTA-FBS. Tissue homogenates were centrifuged at 1500 RPM for 10 min at 4 . Dissociated splenocytes from each mouse were resuspended in 3ml of 1X RBC lysis buffer (containing 0.1M EDTA, 155 mM NH_4_Cl and 10 mM KHCO_3_), incubated for 3 min at RT and washed with PBS-EDTA-FBS buffer. Another centrifugation was performed at 1500g for 10 min at 4 followed by resuspension of cells in 1 ml dissociation buffer. The cell suspension was filtered again using a 100µm mesh filter. Cell counting was performed in a hemocytometer and 2 million cells from each spleen were used for immunostaining and analysis in a flow cytometer (BD LSRFortessa X-20, RRID:SCR_025285).

The antibodies used in the present study are listed in Table 2. Flow cytometry staining was performed at 4°C in canonical bottom 96-well plate (Nunc, Thermo scientific, 249570) protected from light. For all washes, PBS-EDTA-FBS buffer was added, the cells were centrifuged 40s at 1500 rpm, following which the supernatant was discarded, and the cells were re-suspended in the same buffer. To exclude dead cells, the pellet was resuspended in fixable Viability Dye eFluor450 (1:1000) (eBiosciences, Catalog number: 65-0863-18), incubated 15 min and washed. To ensure specific binding with the antibody, the non-specific binding was blocked by using anti-CD16/32 Fc-blocking solution (0.5 ug/millions of cells) for 15 min. After a wash, the pellet was resuspended in a mix of antibodies for 30 min. The cells were washed prior to being fixed for 10 min in 4% PFA. Controls as Fluorescence Minus One (FMO) were included to ensure appropriate identification of a true positive signal and adequate gating strategy. Compensation beads (OneComp eBeads, eBiosciences Inc., San Diego, CA, USA, catalog # 01-1111-42) were incubated in the presence of the fluorescently coupled antibodies to determine the appropriate compensation parameters for each acquisition. Flow cytometry was performed on BD LSRFortessa X-20, BD Biosciences and data were analyzed with FlowJo software (BD Biosciences https://www.flowjo.com/ (RRID:SCR_008520).

**Table 2:** Fluorochrome conjugated antibody panel used for flow cytometry analysis.

| Name of antibody | Species raised in | Dilution used | Manufacturer | Catalog Number/RRID |
| --- | --- | --- | --- | --- |
| CD45-BV711 | Rat | 3ug/mouse | Biolegend | Cat# 103147, RRID:AB_2564383 |
| CD45-FITC | Rat | 1:100 | Biolegend | Cat# 103107, RRID:AB_312972 |
| CD11b-BV605 | Rat | 1:200 | Biolegend | Cat# 101237, RRID:AB_11126744 |
| CD11C-PECY7 | Hamster | 1:200 | BD Biosciences | Cat# 561022, RRID:AB_2033997 |
| LY6C-AF700 | Rat | 1:100 | BD Biosciences | Cat# 561237, RRID:AB_10612017 |
| LY6G-BV510 | Rat | 1:100 | Biolegend | Cat# 127633, RRID : AB_2562937 |
| F4/80-A647 | Rat | 1:200 | Biolegend | Cat# 123121, RRID:AB_893492 |
| CD86-BV650 | Rat | 1:200 | Biolegend | Cat# 105035, RRID:AB_11126147 |
| CD3e-PerCp Cy5.5 | Armenian hamster | 1:200 | Biolegend | Cat# 100328, RRID:AB_893318 |
| CD8a-A647 | Rat | 1:100 | Biolegend | Cat# 100724, RRID:AB_389326 |
| CD4-APC Cy7 | Rat | 1:100 | Biolegend | Cat# 100525, RRID:AB_312726 |
| CD19-BV786 | Rat | 1:100 | BD Biosciences | Cat# 563333, RRID:AB_2738141 |
| NK1.1-BV650 | Mouse | 1:50 | Biolegend | Cat# 108735, RRID:AB_11147949 |

### Bulk RNA extraction and sequencing

RNA extraction from the striatum was performed using the RNAeasy mini kit (Qiagen, 74104) following the manufacturers’ instructions. The quality of the RNA was validated using a BioAnalyzer (Agilent). 600 ng total RNA (RNA integrity number >=7) from each sample was used for sequencing. Briefly, a poly-A selection was performed, followed by an RNA library preparation using KAPA RNA Hyperprep kit (Roche, Kit Code: KK8504). Sequencing was performed with a NextSeq500 (Illumina) instrument using NextSeqHighOutput 75 cycles programme. 20 million single-end 75bp reads were obtained per sample. The output format was saved as Fastq files followed by subsequent DEG analysis.

The resulting FASTQ files were aligned to the mm10 genome reference using the STAR aligner with default parameters (https://github.com/alexdobin/STAR) (RRID:SCR_004463). Raw read counts were quantified using HTSeq-count (https://github.com/htseq/htseq) (RRID:SCR_011867) (75). The DESeq2 package in R (R version 4.3.0, https://www.r-project.org/, RRID:SCR_001905) was utilized for normalization, variance stabilization, and transformation of raw read counts (https://github.com/thelovelab/DESeq2) (RRID:SCR_015687). Differentially expressed genes (DEGs) were identified with DESeq2 (76), applying thresholds of p-value < 0.05, log2 fold change > 1, and an average basal expression > 10. The basal expression threshold ensured the inclusion of actively expressed genes while excluding non-expressed ones.

Principal Component Analysis (PCA) was performed on the normalized read counts of expressed genes (read counts > 10) to explore transcriptome-wide variability among samples. PCA plots were generated using the PCA function in Python (https://www.python.org/downloads/release/python-360/, RRID:SCR_008394). Gene Ontology (GO) analysis of significantly differentially expressed genes was performed with gProfiler (http://biit.cs.ut.ee/gprofiler/ (RRID:SCR_006809) (77, 78). Pathway-level enrichment was assessed using single-sample gene set enrichment analysis (ssGSEA) implemented in GenePattern, (http://www.broadinstitute.org/cancer/software/genepattern) (RRID:SCR_003201) (79). The analysis employed the C5.go.bp.v2022.1.Hs.symbol.gmt database, which includes 7,763 gene sets derived from GO biological processes. Inflammation-related pathways were filtered, and t-tests were conducted on enrichment scores between groups to calculate p-values.

For evaluation of endothelial barrier associated genes, normalized read counts obtained following DESeq2 analysis were extracted for key tight junction markers, including *Ocln*, *Cldn3*, *Cldn5*, *Cldn12*, *Tjp3*, and *Cdh5*. These values were analyzed prior to DEG filtering to assess baseline transcriptional abundance. Statistical comparisons between WT infected and KO infected mice were performed using unpaired t-tests.

### Cytokine analysis in serum and striatal lysates

Serum samples were collected from infected and non-infected mice of both genotypes post MRI at both day 13 and 26 days post-infection. Striatal lysates prepared using RIPA buffer. 30 µg of the striatal lysate from each mouse brain was used to quantify the cyto-chemokines. The cyto-chemokine assay was performed by Eve Technologies (32-plex assay, Calgary, Alberta).

### Statistical analysis

p-values were calculated using GraphPad Prism version 9.4.1,1http://www.graphpad.com/ (RRID:SCR_002798). Statistical differences among the experimental groups were calculated using two-way ANOVA and Tukey’s post-hoc test. Data is represented as mean ± S.E.M.

## Data and materials availability

The data, code, and key lab materials used and generated in this study are listed in a Key Resource Table alongside their persistent identifiers at 10.5281/zenodo.20091139.

## Acknowledgement

This research was funded in part by Aligning Science Across Parkinson’s (ASAP-000525) (to M.D., L.-E.T., H.M., S.G., J.A.S.) through the Michael J. Fox Foundation for Parkinson’s Research (MJFF) and in part by the McGill University Healthy Brains, Healthy Lives (HBHL) grant (to L-E-.T., D.R. and H.M.). The Trudeau lab also receives support from the Canadian Institutes of Health Research (Grant PJT-165928) and from the Krembil Foundation. SM was supported by FRQS (*Fonds de recherche du Québec, Santé*) (2020–2022) and Parkinson Canada (2022–2024) postdoctoral fellowships. The funders had no role in study design, data collection and analysis, decision to publish, or preparation of the manuscript.

We acknowledge the help of the genomics and flow cytometry core facility of the Institut de recherche en immunologie et en cancerologie (IRIC) of the Université de Montréal, with special thanks to Raphaelle Lambert, Annie Gosselin and Angélique Bellemare-Pelletier. We appreciate the technical assistance received from Dr. Charles Ducrot. We also thank Dr. Véronique Laplante and Dr. Lilia Rodriguez for their help in overseeing the data sharing strategy. For the purpose of open access, the author has applied a CC-BY 4.0 public copyright license to all Author Accepted Manuscripts arising from this submission.

An earlier version of this manuscript was posted to BioRxiv on 24/12/2024 at https://www.biorxiv.org/content/10.1101/2024.12.24.630165v1.

## Author contributions

**Conceptualization:** Louis-Eric Trudeau, David A. Rudko, Heidi McBride.

**Data curation:** Sriparna Mukherjee, Vladimir Grouza, Alex Tchung, Amandine Even, Moein Yaqubi, Marius Tuznik, Tyler Canon, Sherilyn Junelle Recinto, Christina Gavino, Marie-Josée Bourque.

**Formal analysis:** Sriparna Mukherjee, Vladimir Grouza, Alex Tchung, Amandine Even, Moein Yaqubi, Nicolas Giguère.

**Funding acquisition:** Louis-Eric Trudeau, David A. Rudko, Samantha Gruenheid, Jo Ann Stratton, Heidi McBride, Michel Desjardins.

**Investigation:** Sriparna Mukherjee, Vladimir Grouza, Alex Tchung, Amandine Even, Moein Yaqubi, Marius Tuznik, Tyler Canon, Sherilyn Junelle Recinto, Christina Gavino, Marie-Josée Bourque.

**Methodology:** Sriparna Mukherjee, Vladimir Grouza, Alex Tchung, Amandine Even, Moein Yaqubi.

**Project administration:** Sriparna Mukherjee, Louis Eric Trudeau

**Resources:** Louis Eric Trudeau, David A Rudko, Samantha Gruenheid, Jo Anne Stratton. **Software:** Alex Tchung, Vladimir Grouza, Moein Yaqubi, Amandine Even, Nicolas Giguère. **Supervision:** Louis Eric Trudeau.

**Validation:** Louis Eric Trudeau.

**Visualization:** Louis Eric Trudeau, Sriparna Mukherjee.

**Writing -original draft preparation:** Sriparna Mukherjee, Vladimir Grouza, Louis-Eric Trudeau.

**Writing- review and editing:** Louis-Eric Trudeau, David A Rudko, Samantha Gruenheid, Jo Anne Stratton.

