## Supplementary information for "A single *Citrobacter rodentium* infection in Pink1 knockout and wildtype mice leads to regional blood-brain-barrier perturbation and limited microglial activation without dopamine neuron axon terminal loss"

***** Denotes equal contribution

**Address correspondence to**:

Dr. Louis-Eric Trudeau

Professor

Faculty of Medicine

Université de Montréal

**S1_Table: List of reagents used for western-blot**

| **Name of Reagent** | **Manufacturer** | **Catalog Number** |
| --- | --- | --- |
| Resolving gel buffer | BioRad | #1610798 |
| Stacking gel buffer | BioRad | #1610799 |
| 10X Tris/Glycine/SDS buffer | BioRad | #1610732 |
| 10X TBS | BioRad | #1706435 |
| Ponceau S solution | Sigma | P7170-1lit |
| 30% Acrylamide/Bis solution 29:1 | BioRad | #1610156 |
| Trans-blot turbo 5X transfer buffer | BioRad | #10026938 |
| TEMED | BioRad | TEM001.50 |
| Ammonium Persulfate | Sigma | A3678-100G |

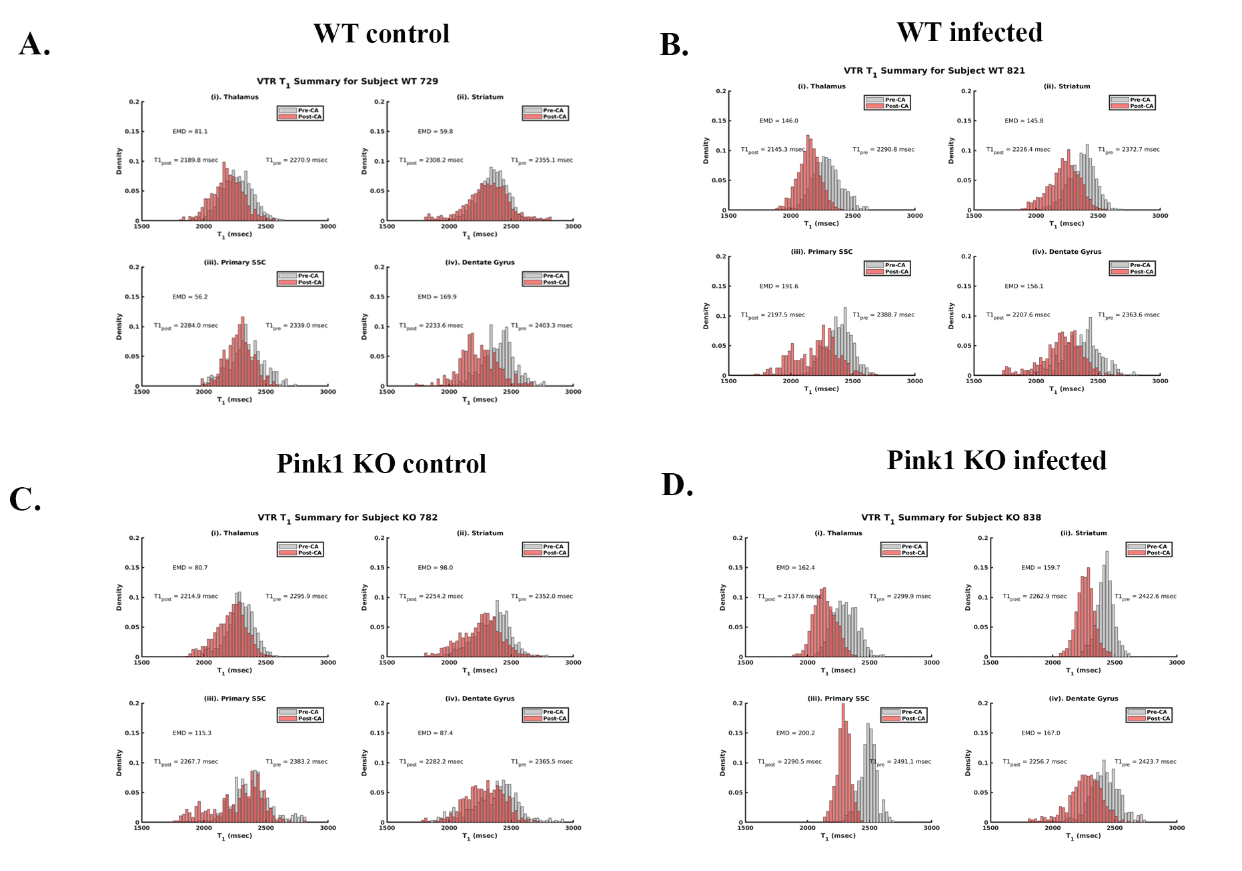

**S1 Fig.** Histograms depicting T1 shift upon gadolinium-based contrast agent injection in the WT and Pink1 KO mice, with or without infection with *Citrobacter rodentium*. The images represent one mouse from each experimental group of two independent cohorts used in the study; WT control in panel (A), WT infected in panel (B), KO control in panel (C) and KO infected in panel (D). Each panel also depicts T1 values (pre-CA as grey and post-CA as red histograms) in four anatomical brain locations namely thalamus (i), striatum (ii), primary SSC (iii) and dentate gyrus (iv).

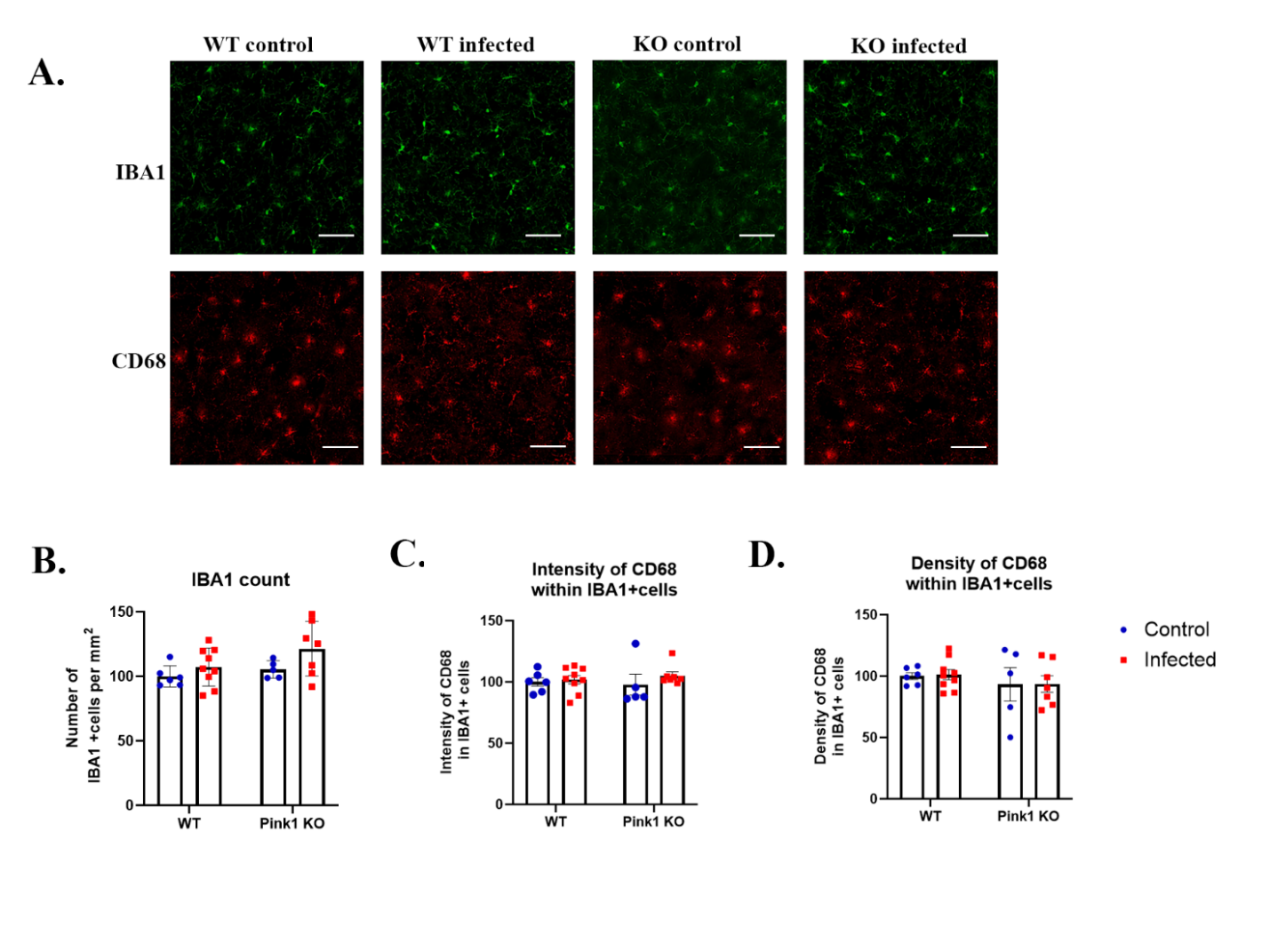

**S2 Fig.** IBA1 and CD68 co-immunostaining in striatal sections from control and infected WT and KO mice. IBA1 was used to identify microglia, and CD68 immunoreactivity was assessed as a marker of lysosomal/phagocytic activation (A) (20X magnification). Quantification includes the number of IBA1+ cells (B) and IBA1+ cell density as well as CD68 intensity and density within IBA1+ microglia (C, D). No significant differences were observed in microglial number or CD68 expression among groups (p > 0.05), indicating an absence of phagocytic engagement following infection. Data are presented as mean ± SEM (scale bar 20µm).

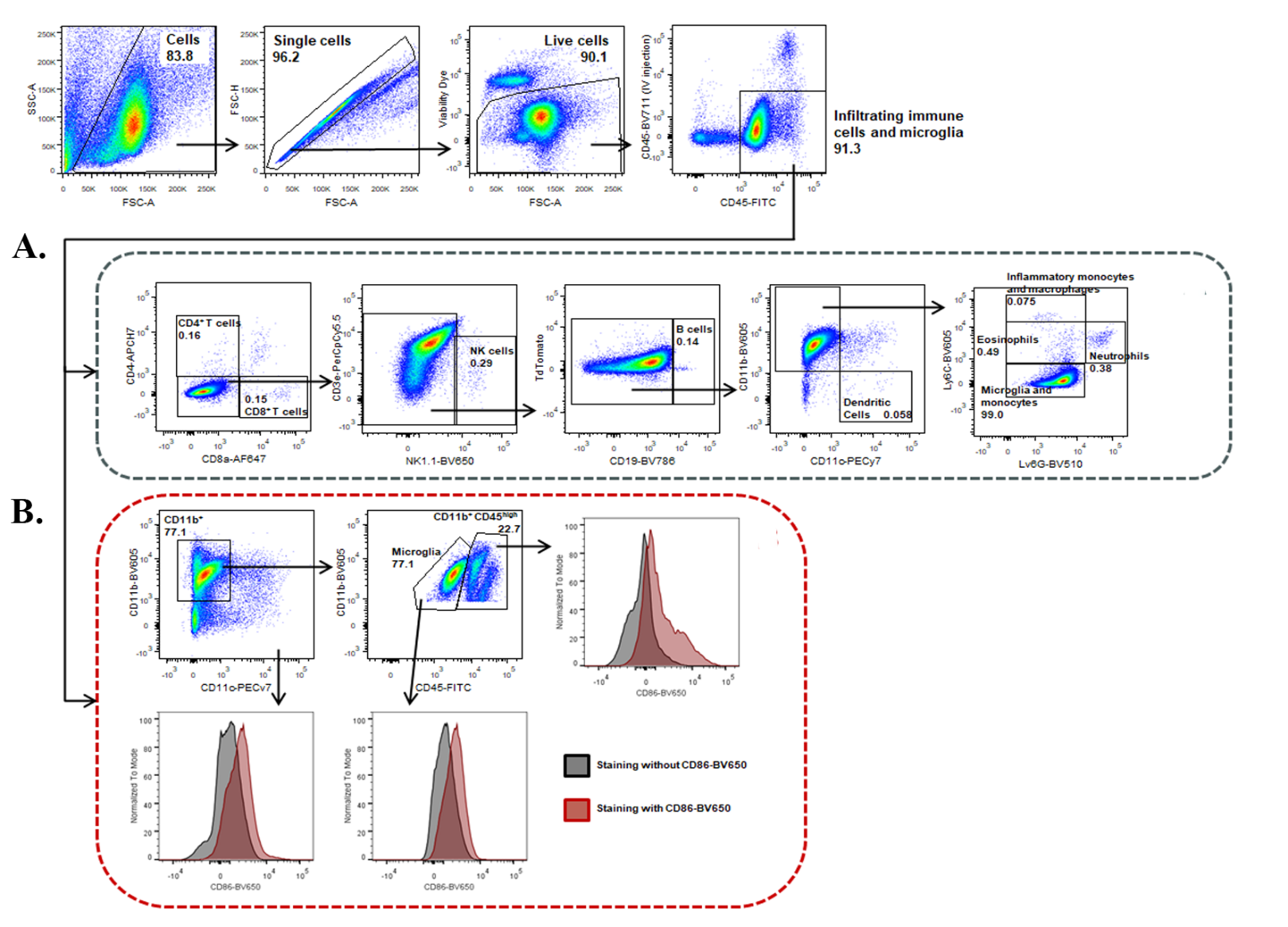

**S3 Fig.** Gating strategy used to determine the percentage and phenotype of immune cell populations in the spleen. Mice were anesthetized and injected with 3 µg of BV711-conjugated anti-CD45 antibody. After collecting and isolating spleen cells, they were stained with various markers. The strategy included the following key steps: Debris and Doublets Exclusion: initially, debris and doublets were excluded from the analysis; Immune Cell Identification: among the viable cells, those with high CD45 expression were identified as immune cells. Within this CD45high gate, NK cells were identified as CD3-NK1.1+, CD4+ T cells were CD3+CD4+CD8-, CD8+ T cells were CD3+CD4-CD8+, B Cells were CD19+, dendritic cells were CD11c+CD11b, monocytes were CD11b+Ly6C- Ly6G-, neutrophils were CD11b+Ly6C^int^ Ly6G+, eosinophils were CD11b+Ly6C^int^ Ly6G- and inflammatory monocytes and macrophages were CD11b+Ly6C^high^ Ly6G- (A). Within immune cells, the expression of CD86 was analysed in CD11b-CD11c+ dendritic cells, CD11b+F4/80-Ly6C- macrophages, CD11b+F4/80-Ly6C-Ly6G- monocytes, CD11b+F4/80-Ly6C^int^Ly6G+ neutrophils, CD11b+F4/80-Ly6C^int^Ly6G^-^eosinophils and CD11b+F4/80-Ly6C^high^Ly6G- inflammatory monocytes. The red histograms display CD86 expression for these cell populations, while the black histograms represent fluorescence minus one (FMO) control, excluding the CD86 antibody. The plot and histograms are representative of data from three independent experiments.

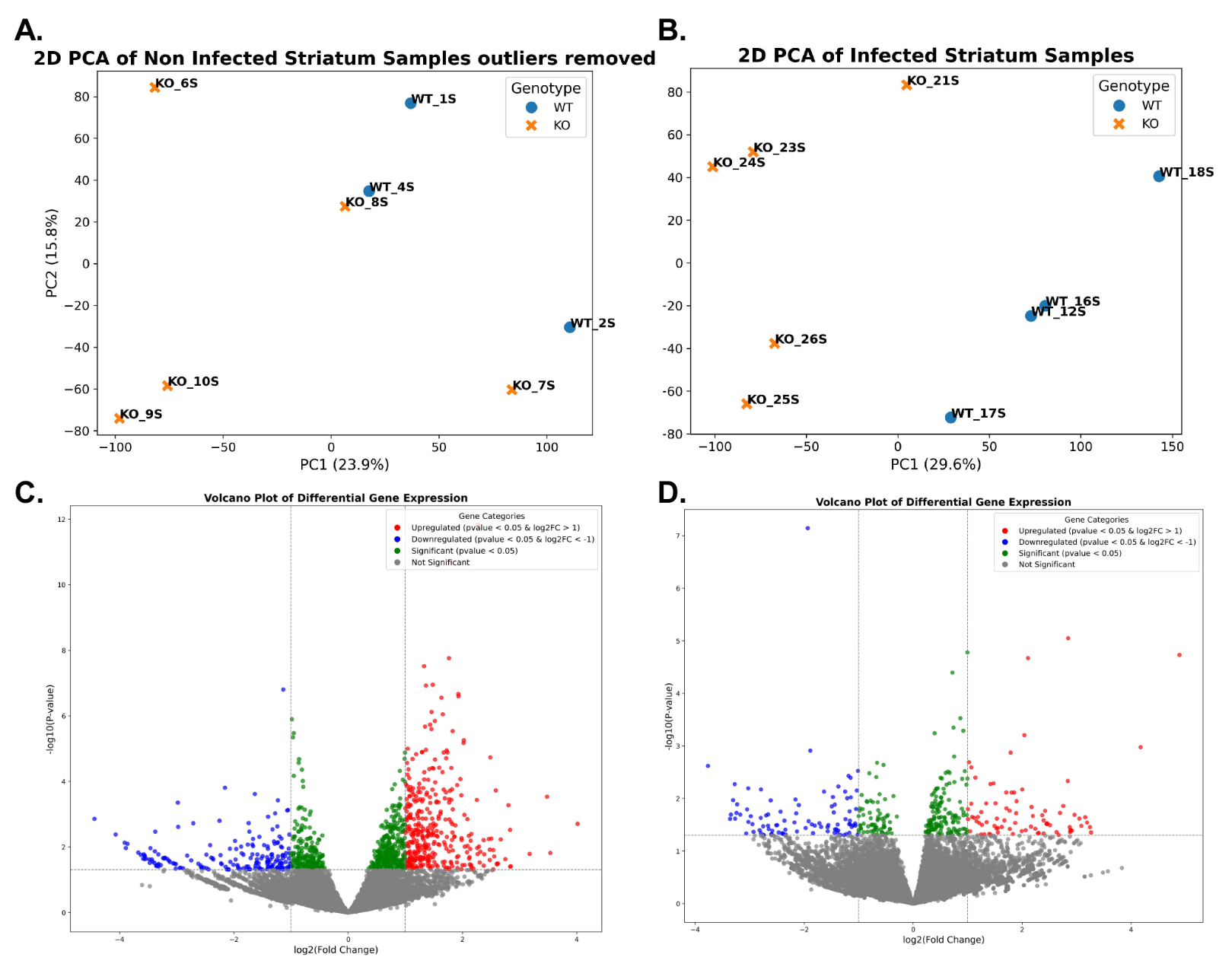

PC2(12.3%)

**S3 Fig.** Principal Component Analysis (PCA) and Volcano plots of the differential gene analysis of the striatal transcriptome data. PCA plot showing the distribution of transcriptomic data for samples across Principal Component 1 (PC1) and Principal Component 2 (PC2), which explain 23.9% and 15.8% of the total variance of the non-infected and 29.6% and 12.3% of the infected samples respectively (A, B). Each point represents an individual sample, color-coded by treatment group (WT in blue, Pink1 KO in orange). Clustering of samples reflects similarity in their gene expression profiles. Outlier samples, which did not cluster with others in the same treatment group, were identified and excluded from subsequent analyses. The volcano plot visualizes the results of differential gene expression analysis between WT and KO for both non-infected and infected conditions (C, D). Each point represents an individual gene, plotted according to its log2 fold change (x-axis) and -log10 adjusted p-value (y-axis). Genes upregulated are highlighted in red, while downregulated genes are shown in blue. Non-significant genes are displayed in gray and the significant ones among both up and downregulated genes are highlighted in green for both non-infected and infected groups.

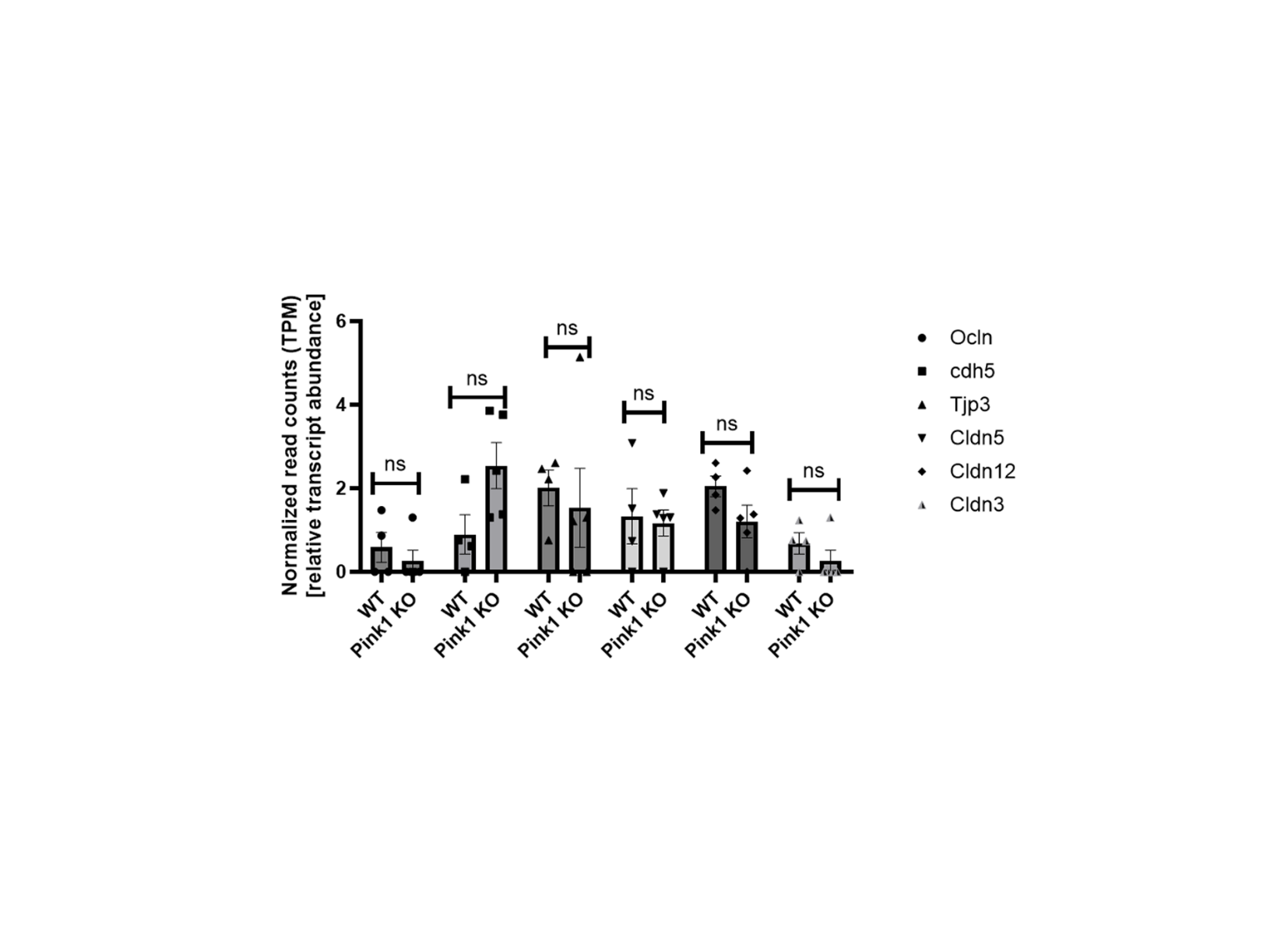

**S6 Fig.** Expression of tight junction related genes in infected WT and Pink1 KO mouse striatum. Normalized RNA-seq read counts (TPM, relative transcript abundance) for canonical BBB junctional markers including Ocln, Cdh5 (VE-cadherin), Tjp3, Cldn5, Cldn12, and Cldn3 in striatal tissue from infected WT and Pink1 KO mice at day 26 post infection. Individual data points represent values from single animal (WT, n =4; KO, n = 5), with bars indicating mean ± SEM. Statistical comparisons between infected WT and KO groups were performed using unpaired t-tests with Welch’s correction; no significant differences were detected for any of the genes (all p > 0.05).
